# VAP-mediated membrane tethering mechanisms implicate ER-PM contact function in pH homeostasis

**DOI:** 10.1101/2023.01.12.523872

**Authors:** Kar Ling Hoh, Baicong Mu, Tingyi See, Amanda Yunn Ee Ng, Annabel Qi En Ng, Dan Zhang

**Author notes:** Address correspondence to Dan Zhang. These authors contributed equally to this work (Kar Ling Hoh, Baicong Mu, Tingyi See).

## Abstract

VAMP-associated proteins (VAPs) are highly conserved endoplasmic reticulum (ER) resident proteins that establish ER contacts with multiple membrane compartments in many eukaryotes. However, VAP-mediated membrane tethering mechanisms remain ambiguous. Here, focusing on fission yeast ER-plasma membrane (PM) contact formation, using systematic interactome analyses and quantitative microscopy, we predict a non-VAP-protein direct binding-based tethering mechanism of VAPs. We further demonstrate that VAP-anionic phospholipids interactions underlie ER-PM association and define the pH-responsive nature of VAP-tethered membrane contacts. Importantly, such conserved interactions with anionic phospholipids are generally defective in amyotrophic lateral sclerosis (ALS)-associated human VAPB mutant. Moreover, we identify a conserved FFAT-like motif locating at the autoinhibitory hotspot of the essential PM proton pump Pma1. This modulatory VAP-Pma1 interaction is crucial for pH homeostasis. We thus propose an ingenious strategy for maintaining intracellular pH by coupling Pma1 modulation with pH-sensory ER-PM contacts via VAP-mediated interactions.

## Introduction

In eukaryotes, membrane-bound organelles with characteristic morphologies, dynamics, and functions, are interconnected at membrane contact sites (MCSs) where respective activities can be jointly regulated. A number of conserved tethering proteins has been identified at different MCSs ^1, 2^. Among them, VAPs (Scs2 and Scs22 in yeasts; VAPA and VAPB in human) are the only ER resident anchors connecting various proteins at multiple MCSs via FFAT (two phenylalanines/F in an acidic tract) or its related motifs. Each VAP contains an N-terminal conserved major sperm protein (MSP) domain bathing in the cytosol and a transmembrane segment at the extreme C-terminus ^3, 4^. In both budding and fission yeasts, VAPs are the major tethers for ER-PM association ^5, 6^. Unlike other yeast ER-PM tethering proteins, namely tricalbins (Tcb1/2/3) and Ist2 which directly bind acidic phospholipids (PLs) in the PM ^7, 8^, molecular basis for VAP-mediated ER-PM contact formation is however still unknown.

VAP-mediated ER-PM contacts have been functionally linked to lipid homeostasis in yeast. VAPs recruit OSBP-related protein (ORP)/oxysterol-binding homology (Osh) family via FFAT-binding to ER-PM junctions and regulate their phosphatidylinositol 4-phosphate (PI_4_P) /sterol transferring activities to keep PI_4_P level in the PM ^9^. Other roles of ER-PM contacts in ER organization and stress signaling have also been suggested ^6^. Moreover, tight ER-PM junctions can impose steric effects on cortical events. Such physical roles have been mainly established for contractile ring assembly and for delimiting exocytic territory in the fission yeast *Schizosaccharomyces pombe* (*S. pombe*) ^10–12^. Here, for the first time, we expand VAP-mediated ER-PM contact functions to pH homeostasis using *S. pombe*.

In the PM of fungi and plant cells, proton pumping P-type adenosine triphosphatases (ATPase) (known as Pma1 in yeast) maintain the intracellular pH and membrane potential essential for nutrient uptake ^13^. Pma1 assembles as hexamers ^14^. Each monomer contains multiple transmembrane _α_-helices (TMs) and four major cytosolic domains: the actuator (A) domain, the nucleotide-binding (N) domain, the phosphorylation (P) domain, and the C-terminal regulatory (R) domain which is autoinhibitory by locking other cytosolic domains via intra- or inter-molecular interactions ^15, 16^. Glucose-dependent phosphorylation of R domain has been postulated to activate these pumps in yeast by abolishing such interactions^17^. However, it is unknown if other glucose-independent mechanisms have been evolved for Pma1 regulation.

In this study, we provided first *in vivo* evidence to demonstrate that VAPs establish pH-sensitive ER-PM contacts through interactions with major PM anionic PLs, especially high-abundance phosphatidylinositol (PI) and phosphatidylserine (PS). Our finding thus recognizes the MSP as a conserved lipid-binding domain and unifies that lipid-protein interactions dominate ER-PM contact formation. Our interactome analyses also revealed major VAP interactors and implicated potential roles of VAPs in various cellular events. Importantly, we pinpointed a conserved FFAT-like motif in Pma1 at its autoinhibitory P-R domain binding site and confirmed that such VAP interaction is protective against cytoplasmic acidification and crucial for pH homeostasis. Our study hence indicates a new mechanism for Pma1 regulation by hijacking its autoinhibition via competitive VAP-P domain binding. Collectively, we propose that VAPs sustain cytosolic pH by finetuning their interactions between Pma1 and pH-biosensory PLs at ER-PM contacts.

## Results

### VAPs form ER-PM contacts via MSP domain in a dosage-dependent manner

A previous study has shown that Scs2 plays a dominant role in ER-PM tethering over Scs22 in fission yeast ^5^ (see also Fig. 1a). In fact, Scs2 was expressed at a much higher level than Scs22 in wild type (*WT*) (Fig. 1b). Interestingly, ER-PM contact formation was also compromised when we swapped the open reading frames of *scs2* and *scs22* at their genomic loci (Supplementary Fig. 1a), implying that there might be intrinsic differences between two VAPs in their ER-PM tethering function.

**Figure 1.**
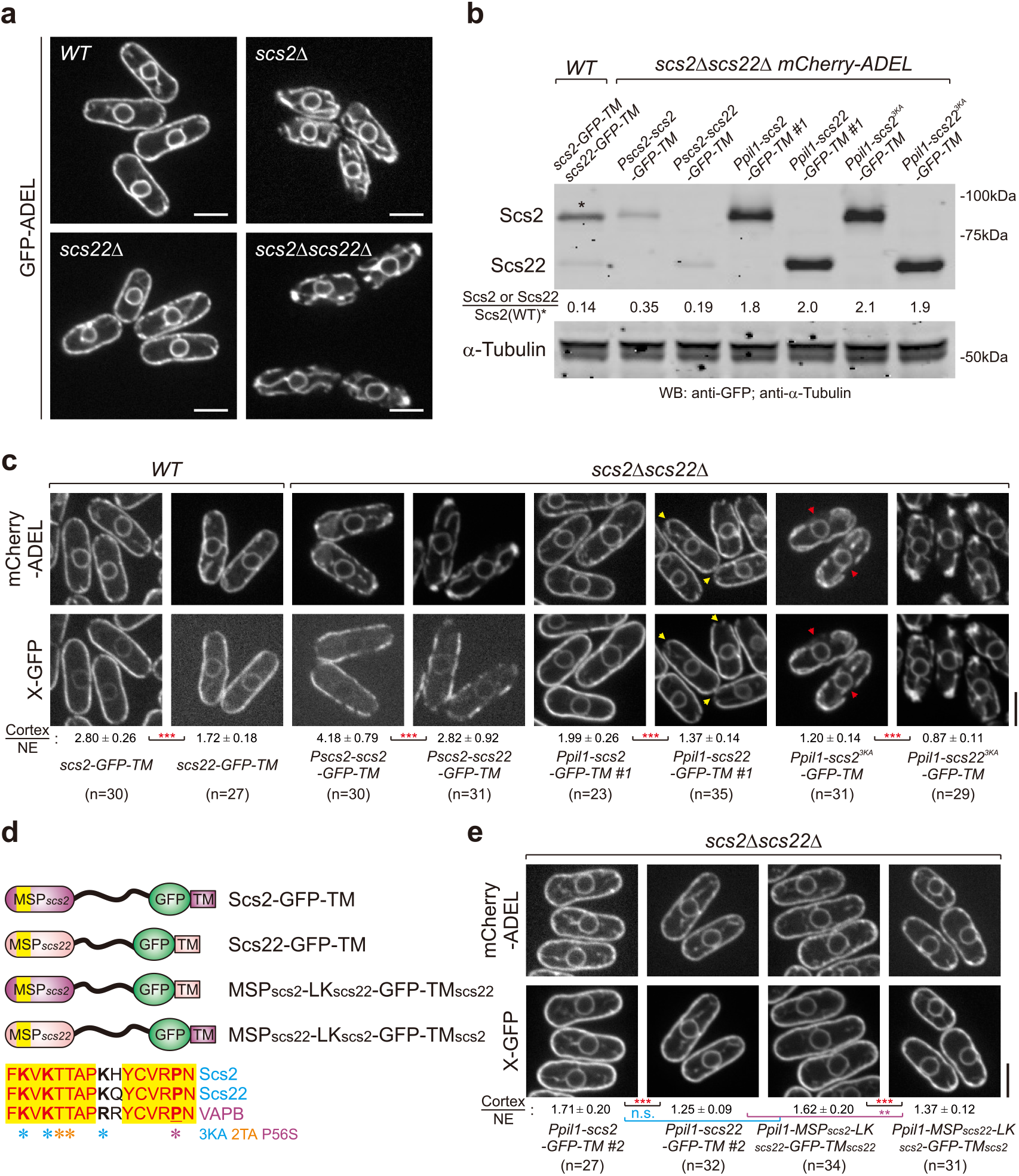
Fission yeast VAPs mediate ER-PM contact formation through the conserved MSP domain in a dosage dependent manner. (**a**,**c**,**e**) Spinning disk confocal images of indicated cells expressing indicated proteins. Shown are central focal planes. Yellow arrowheads point to cell tips lacking ER-PM contacts. Red arrowheads point to lateral cortical regions lacking ER-PM contacts. Normalized cortex/NE ratios (mean±standard deviation [SD]) of fission yeast VAP variants are included. *p*-values, two-tailed t test, \*\*\**p*<10^−7^<\*\**p*<0.0001; n.s., not significant. n, cell number. Scale bars, 5 μm. (**b**) Expression levels of GFP-tagged VAP variants in indicated cells. Relative ratios of protein levels as compared to Scs2 level (asterisked) in wild type (*WT*) are included. Protein samples were probed with anti-GFP and anti–α-Tubulin antibodies. (**d**) The cartoon shows the design of VAP swop variants, and the alignment of consensus sequences from fission yeast VAPs and human VAPB. TM, the C-terminal transmembrane domain. Blue and orange asterisks show mutated sites in this work and the magenta asterisk indicates the ALS-associated VAPB mutation.

Unlike Scs22 where its nuclear envelope (NE) localization was obvious, the NE pool of Scs2 was barely visible, suggesting likely stronger affinity of Scs2 towards the cell cortex (Fig. 1c). We then ectopically expressed Scs2 or Scs22 at various levels in cells lacking native VAPs (Fig. 1b). We chose the promoter of *pil1* that encodes an abundant eisosome subcomponent ^18^ to achieve high protein expression. Scs2 always exhibited higher cortex-to-NE ratios than Scs22 under similar expression levels. Moreover, despite both VAPs could dose-dependently restore ER-PM contacts in *scs2Δscs22Δ* cells, Scs2 appeared more competent in ER-PM tethering (Fig. 1c). We envisaged that strong PM binding of an ER-PM tether might dampen its mobility. In line with this, the artificial tether TM-GFP-CSS_Ist2_ which resides in the ER and binds to the PM ^5^ displayed a slower fluorescence recovery after photobleaching (FRAP) along the cell cortex than the control ER-resident TM-GFP (Supplementary Fig. 1b). Similarly, Scs2 always diffused slower than Scs22 in *WT* or in *scs2Δscs22Δ* cells when expressed in comparable levels (Supplementary Fig. 1c). We thus conclude that Scs2 harbors an intrinsic higher PM affinity than Scs22.

Notably, mutations of three lysine residues (K36/38/43A, hereafter referred to as 3KA) in the VAP consensus sequence (VCS) within the MSP domain ^19^ of either Scs2 or Scs22 abolished their ER-PM tethering capacity regardless of high expression levels (Fig. 1b-d). As expected, both Scs2^3KA^ and Scs22^3KA^ nearly lost the PM affinity, manifested by either reduced cortex-to-NE ratios or increased mobility to levels of non-tether ER proteins (i.e., the ER luminal marker mCherry-ADEL ^20^ for cortex-to-NE ratios and TM-GFP for mobility respectively in Fig. 1c and Supplementary Fig. 1c). These data indicate that the PM affinity of a VAP is likely determined by its conserved MSP domain. We then constructed chimeric variants by swopping MSP domains between Scs2 and Scs22 (Fig. 1d) and expressed them in *scs2Δscs22Δ* cells at comparable levels (Supplementary Fig. 1d). Indeed, MSP_Scs2_-containing constructs always exhibited higher cortex-to-NE ratios than the ones with MSP_Scs22_ (Fig. 1e).

Collectively, our results suggest that the conserved MSP domain intrinsically defines dose-dependent ER-PM tethering capacity of VAPs.

### VAPs keep the PM PI_4_P level via MSP domain in a dosage-dependent manner

It has been shown that fission yeast cells lacking both VAPs display elevated PM PI_4_P levels marked by the pleckstrin homology (PH) domain of budding yeast Osh2 ^5^ (see also Supplementary Fig. 1e). Such abnormally high PI_4_P levels of the PM cannot be rescued by artificially re-built ER-PM contacts ^5^, indicating specific roles of VAPs in the PM PI_4_P homeostasis.

As compared to *WT*, we found that levels of the PM PI_4_P remained high in *scs2Δscs22Δ* expressing low levels of either VAPs where ER-PM contact re-establishment was incomplete (Supplementary Fig. 1f). Intriguingly, unlike wild-type VAPs or even chimeric variants that still contained intact MSP domains, high expression of either Scs2^3KA^ or Scs22^3KA^ in *scs2Δscs22Δ* failed to reduce the high PM level of PI_4_P (Supplementary Fig. 1f). These data suggest the necessity of a functional MSP domain for PI_4_P-related activity of VAPs. Therefore, we conclude that roles of VAPs in the PM PI_4_P homeostasis require the MSP domain in a dose-dependent manner.

### Two fission yeast VAPs show divergent binding preferences

We next attempted to explore molecular interactions underlying VAP-mediated membrane tethering. Given that Scs2 and Scs22 showed distinct PM affinities, we speculated that they may also differ in their potential cortical binding partners. We performed interactome analyses using both full-length and TM-truncated VAPs (namely Scs2N and Scs22N) as baits, considering that the TM-truncated fragments are likely more accessible to potential hidden interactors than ER-anchored native VAPs.

Interestingly, many interactors shared by all four baits were lipid enzymes or lipid transporters (Fig. 2a and Supplementary Data 1). Interactors involved in cell polarity or morphogenesis that are localized to cell tips showed an overall binding preference to Scs2 and Scs2N (Supplementary Fig. 2a). This may partly explain the fact that Scs22 was less capable of establishing ER-PM contacts at cell ends (arrowed in Fig. 1c). Proteins that organize the lateral cell cortex were also detected to preferentially bind Scs2 (Supplementary Fig. 2a). Notably, 49 out of total 296 interactors detected with at least one bait were related to vesicle-mediated trafficking. Among these major hits, regulatory proteins mainly associated with Scs22, while coatomers and SNARE/NSF-related factors appeared to favor Scs2 binding (Supplementary Fig. 2a). We then picked the top five Scs2-favored interactors which localized to the lateral PM and verified such interactions using Co-immunoprecipitation (Co-IP) (Fig. 2b and Supplementary Fig. 2b) ^21^. We also confirmed Scs2-Scs22 interaction by Co-IP (Supplementary Fig. 2c).

**Figure 2.**
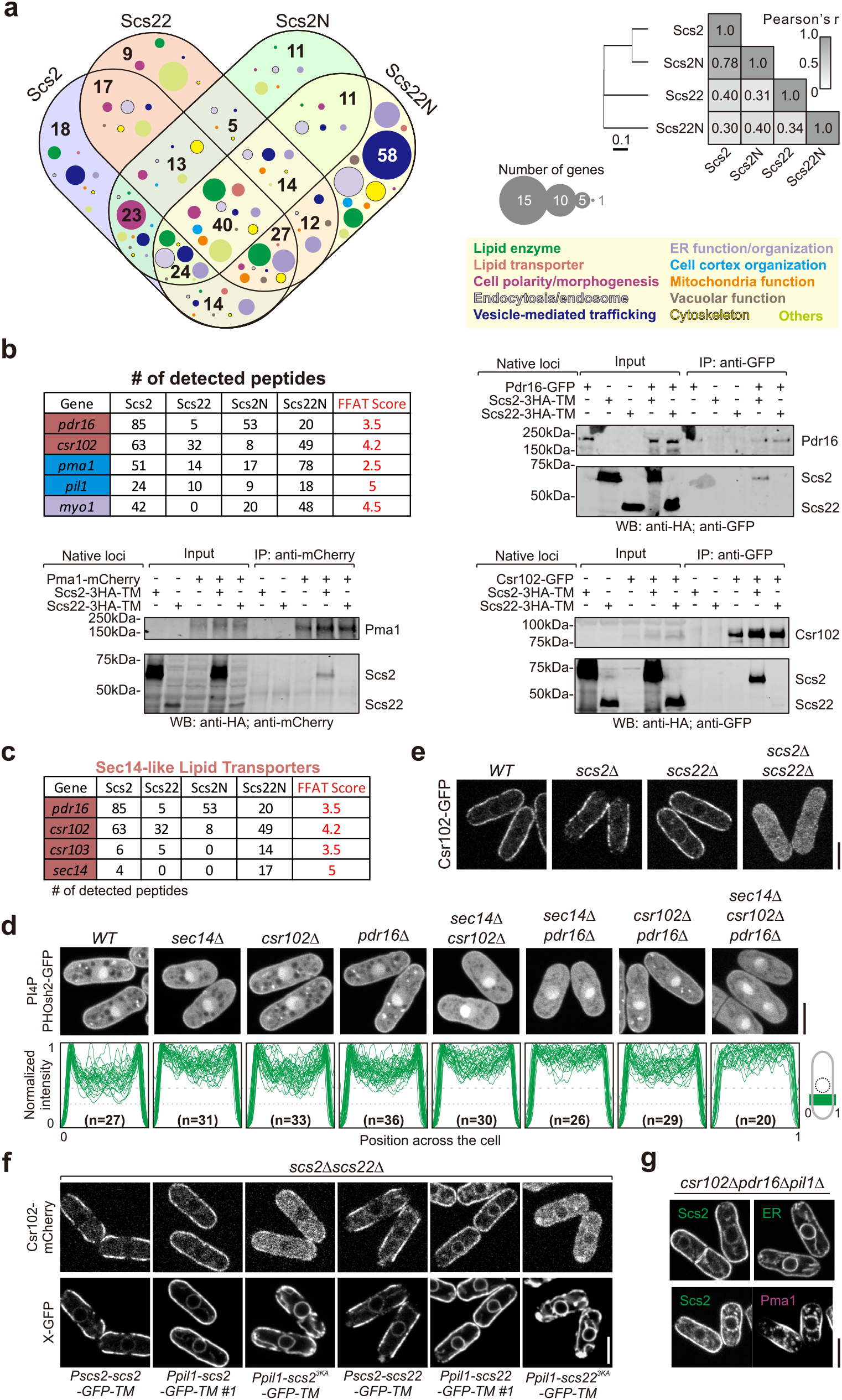
Interactome analyses implicate VAPs in functional modulation of Sec14 family and predict a non-VAP-protein direct interaction-based ER-PM tethering in fission yeast. (**a**) A statistical overview of interactome results summarized from Supplementary Data 1. Numbers of overlapped interactors are included in the Venn diagram. Scale bar, cophenetic distance. (**b**) Co-IP of indicated proteins expressed from native loci. Samples were probed with indicated antibodies. The top five hits of Scs2 interactors localizing to the lateral cell cortex are shown in the table. (**c**) Sec14-like lipid transporters detected in interactome analyses are shown with numbers of detected peptides and the corresponding minimal FFAT scores. (**d-g**) Scanning confocal images of indicated cells expressing indicated proteins. Shown are central focal planes. In (**d**), intensity profiles of PH_Osh2_-GFP over illustrated regions of analyzed cells were shown in the bottom. n, cell number. Scs2, Scs2-GFP-TM; ER, the luminal marker GFP-ADEL. Pma1, Pma1-mCherry. Scale bars, 5 μm.

Many VAP-interacting proteins contain the FFAT motif which has been shown to bind the MSP domain via electrostatic interactions ^22^. A position weight matrix has been established for calculating a FFAT score of any given 13-residue motif that predicts its VAP binding potential (FFAT score of an ideal motif is 0 and 2.5 is the typical cutoff score) ^4, 23^. Remarkably, most interactors we detected or confirmed did not contain a typical FFAT motif (Fig. 2b, Supplementary Fig. 2a-d and Supplementary Data 1). We also did not detect peptide-enrichment of proteins with low FFAT scores (Supplementary Fig. 2d), implying a possible existence of more complex interaction patterns for fission yeast VAPs beyond FFAT-based association.

Moreover, we noticed that four out of five Sec14-like lipid transporters were detected in Scs2 interactome (Fig. 2c), with previously uncharacterized Csr102 and Pdr16 as top two hits (Fig. 2b). Sec14 family proteins have been suggested to engage in PI transfer, however importance of such lipid-related function *in vivo* is still obscure for both fission and budding yeasts ^24, 25^. We found that both the PM and Golgi pools of PI_4_P decreased significantly in tubby-shaped *sec14Δ* cells. Such reduction was enhanced when cells further lost either or both of Csr102 and Pdr16 in the background, whereas mutants with single or double deletion of *csr102* and *pdr16* remained *WT*-like (Fig. 2d). No detectable change of phosphatidylinositol 4,5-bisphosphate (PI_4,5_P_2_) levels was seen in all these mutants (data not shown). These data suggest that Sec14 family proteins contribute to PI_4_P homeostasis in *S. pombe*.

Interestingly, Csr102 bound both VAPs at their respective native levels, and exhibited a likely higher affinity to Scs22 when VAPs were ectopically expressed in a comparable level in *scs2Δscs22Δ* background (Fig. 2b and Supplementary Fig. 2e). Such interactions were required for its cortical localization, as Csr102 became cytosolic in absence of both VAPs even when ER-PM contacts were artificially restored (Fig. 2e and Supplementary Fig. 2f). Indeed, both Scs2^3KA^ and Scs22^3KA^ that mostly lost Csr102-interaction failed to recruit Csr102 to the cortex (Supplementary Fig. 2e and Fig. 2f). In contrast, the cortical localization of Pdr16 was VAP-independent (Supplementary Fig. 2g). Except Scs2, Pdr16 did not interact with Scs2^3KA^, Scs22 or Scs22^3KA^, even when these VAP variants were highly expressed (Supplementary Fig. 2h and Fig. 2b). Though it is unclear how VAP-interactions would affect Sec14 family activities in PI_4_P homeostasis, it is attractive to speculate such a modulatory role of VAPs at least via Csr102.

In summary, we revealed that Scs2 and Scs22 differed in protein binding preference, especially with cortical proteins.

### A prediction of non-VAP-protein direct binding-based ER-PM coupling

We think that cross-membrane interactions are the basis of establishing MCSs of any kind, including VAP-mediated ER-PM tethering. Dosage-dependent ER-PM tethering of VAPs we hitherto demonstrated somewhat implies the existence of collectively abundant PM interaction partners which could afford unsaturated interactions with VAPs of high expression levels.

To further explore molecular basis of VAP-mediated ER-PM tethering, we focused on the top five Scs2-interactors localizing to the lateral PM (Fig. 2b and Supplementary Fig. 2b). Among them, Myo1 appears least likely a key tethering partner, as it is dynamically associated with endocytic sites and mainly enriched at growing tips with a limited pool along the interphase lateral cell cortex ^26^. To the contrary, Pil1 and the essential H+-ATPase Pma1 are known highly abundant and static landmarks laterally covering distinct PM domains ^18, 27^ (Supplementary Fig. 2i). However, no reduction of ER-PM contacts was seen in *csr102Δpdr16Δpil1Δ* cells, and Scs2 could still establish ER-PM contacts at cortical regions devoid of Pma1 in the background (Fig. 2g). Besides, Scs2 diffused much faster than these PM interactors (Supplementary Fig. 2i), but at similar rates as the artificial ER-PM tether (Supplementary Fig. 1b,c) that binds anionic PLs of the PM ^5, 28^. Furthermore, despite carrying mutations (T39/40A) which impair FFAT-binding in budding yeast ^19^, the highly expressed Scs2^2TA^ successfully restored ER-PM contacts in *scs2Δscs22Δ* cells (Supplementary Fig. 3a), implying that FFAT-interaction is nonessential for VAP-mediated ER-PM tethering. These results also suggest the presence of other major cortical partners and a likelihood of non-VAP-protein direct interaction-based tethering for VAPs.

In fact, we detected rather limited number of ER resident interactors using both full-length Scs2 and cytosolic Scs2N (Supplementary Data 1). We reason that if PM proteins were indeed required for Scs2-mediated ER-PM contact formation, PM-anchored Scs2 would be otherwise an incompetent tether. We thus constructed PM-anchored Scs2 (hereafter referred as Scs2-PM) by replacing its C-terminal TM domain with a PM targeting motif ^29^ and ectopically expressed it under the strong thiamine (Thi)-repressible *nmt1* or *pi11* promoter in *scs2Δscs22Δ* cells. Fascinatingly, though not as fully efficient as *WT*, Scs2-PM was able to establish ER-PM contacts in a dose-dependent manner, which similarly required a functional MSP domain (Fig. 3a). Surprisingly, like *WT* (Supplementary Fig. 1f), Scs2-PM was able to restore the PM PI_4_P level in *scs2Δscs22Δ* cells, while Scs2^3KA^-PM again lost the competence (Fig. 3b). Likewise, the cortical recruitment of Csr102 was fully rescued by Scs2-PM, but not Scs2^3KA^-PM (Supplementary Fig. 3b).

**Figure 3.**
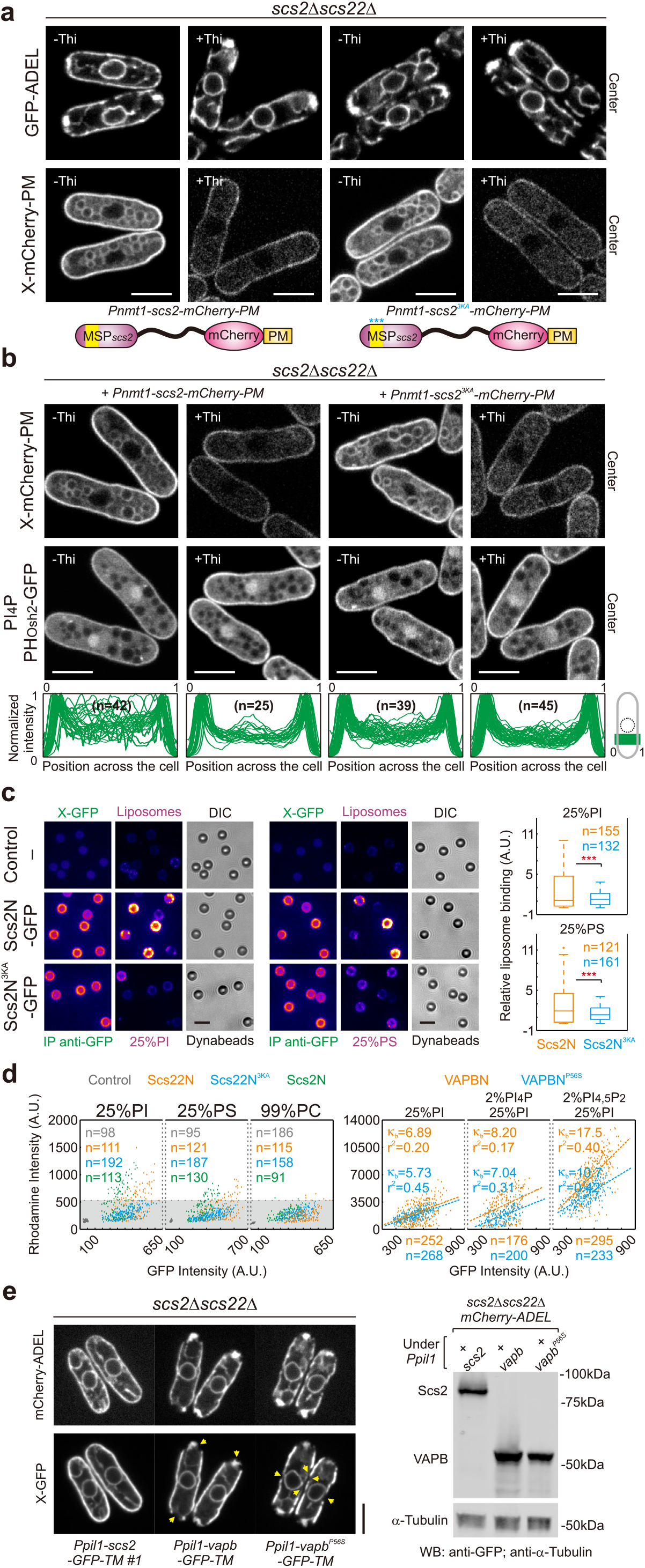
Conserved binding of VAPs with major anionic phospholipids underlies the membrane tethering function. (**a,b**) Scanning confocal images of indicated cells expressing indicated proteins. Shown are central focal planes. The cartoon in (**a**) shows the design of the PM-anchored mCherry-tagged Scs2 variants. In (**b**), intensity profiles of PH_Osh2_-GFP over illustrated regions of analyzed cells were shown in the bottom. Thi, thiamine. n, cell number. (**c,d**) Fluorescence microscopy-based protein-liposome binding assays. Indicated GFP-tagged VAP variants immobilized on Dynabeads were incubated with indicated rhodamine-labelled liposomes (see Methods for details). In (**c**), shown are pseudo-colored spinning disk confocal images with the same contrast for each fluorescent channel. Quantifications are shown on the right. Plots from raw intensity data are included in (**d**); the dotted line and shadow mark the liposome binding threshold. *p*-values, two-tailed t test, \*\*\**p*<10^−5^; κ_b_, liposome binding coefficient; r^2^, r squared; n, bead number. (**e**) Central focal plane scanning confocal images of indicated cells. Yellow arrowheads point to the tip or lateral cortical regions lacking ER-PM contacts. Expression levels of GFP-tagged Scs2 and VAPB variants are shown on the right. Samples were probed with indicated antibodies. Scale bars, 5 μm.

Taken together, our data predict that direct VAP-protein interactions are unlikely the major molecular forces driving Scs2-mediated ER-PM association in fission yeast.

### VAPs interact with major anionic phospholipids

We next sought to explore possible PLs**-**interaction of fission yeast VAPs and envisaged that such potential PLs should be abundant in both the PM and the ER, for instance, main anionic PLs **-** PI and PS ^30, 31^.

In fact, purified Scs2N-6His could bind both PI and PS liposomes. Such binding was enhanced with increased PI or PS concentration, or by adding PI_4_P to respective liposomes (Supplementary Fig. 3c). Unfortunately, recombinant Scs2N^3KA^-6His was unsuitable for liposome co-sedimentation assays as it generally precipitated after centrifugation (data not shown). We thus developed a quantitative fluorescence microscopy-based protein-liposome binding assay (see Methods). In brief, we examined protein-liposome binding using rhodamine-labelled liposomes with magnetic beads that were immuno-conjugated with GFP-tagged cytosolic portions of VAP variants. Concordantly, Scs2N bound specifically to PI and PS, Scs2N^2TA^ showed weaker binding to both, whereas Scs2N^3KA^ exhibited significantly lower affinities for both (Fig. 3c and Supplementary Fig. 3d,e). The PLs-binding capacities of Scs2N variants showed a clear correlation with their PM affinities as indicated by the respective cortex-to-NE ratios (Supplementary Fig. 3a). Likewise, addition of either PI_4_P or PI_4,5_P_2_ to PI liposomes increased Scs2N binding affinity, and such effects were slightly more pronounced with PI_4_P (Supplementary Fig. 3f). Remarkably, Scs22N could also bind to PI and PS, but exhibiting generally weaker binding as compared to Scs2N. Unlike Scs2N^3KA^, Scs22N^3KA^ fully lost binding to PI or PS (Fig. 3d). Notably, such PLs**-**interaction patterns of fission yeast VAPs and their mutants were well correlated with their respective ER-PM tethering capacities as we described above.

We then wondered if such PLs**-**binding capacity is conserved with human VAPB. Interestingly, TM-truncated VAPB (VAPBN) could also bind PI and PS, albeit exhibiting much weaker affinities than Scs2N (Supplementary Fig. 3g). These data may explain the fact that highly expressed VAPB was unable to establish ER-PM contacts near fission yeast cell tips where PS is accumulated ^32^ (Fig. 3e). PI-VAPBN binding was greatly enhanced by PI_4,5_P_2_, while PI_4_P had a minor effect (Fig. 3d). To obtain physiological implications of PI/PIPs-VAPB interactions, we turned to familial ALS8 mutant VAPB^P56S^ ^33^ which harbors a single mutation within the VCS (Fig. 1d). VAPBN^P56S^ showed a comparable PI or PI_4_P binding affinity as VAPBN, but a clearly weaker PI_4,5_P_2_ or PS association (Supplementary Fig. 3g and Fig. 3d). As expected, VAPB^P56S^ was compromised in ER-PM tethering, both at cell tips and along cell sides (Fig. 3e).

In summary, fission yeast VAPs interact with major anionic PLs, which requires the MSP domain. Binding of PI, PS and phosphoinositides appears to be a conserved feature of VAP family.

### VAP-anionic PLs interactions underlie ER-PM tethering mechanism

Given that binding affinity of VAP variants with major anionic PLs was correlative to their ER-PM tethering capacity, we hypothesized that such lipid-protein interactions could be the basis for ER-PM contact formation in fission yeast.

To directly test this idea, we used *pps1Δ* cells lacking the only PS-synthesizing cytidine diphosphate-diacylglycerol (CDP-DAG)-serine O-phosphatidyltransferase in *S. pombe* (Fig. 4a). It has been shown that *pps1Δ* cells have no detectable PS but exhibit an increased PI level ^34^. Indeed, as previously shown ^32^, the PM pool of PS marked by GFP-Lact-C2 ^35^ was absent in *pps1Δ* (Supplementary Fig. 4a). We also saw augmented PM levels of PI_4_P and PI_4,5_P_2_ indicated by PH_Osh2_-GFP and PH_Num1_-GFP ^5^ respectively in these cells (Supplementary Fig. 4a), mirroring overall high PI content. ER-PM contact formation was however unaffected in *pps1Δ* (Fig. 4b and Supplementary Fig. 4b).

**Figure 4.**
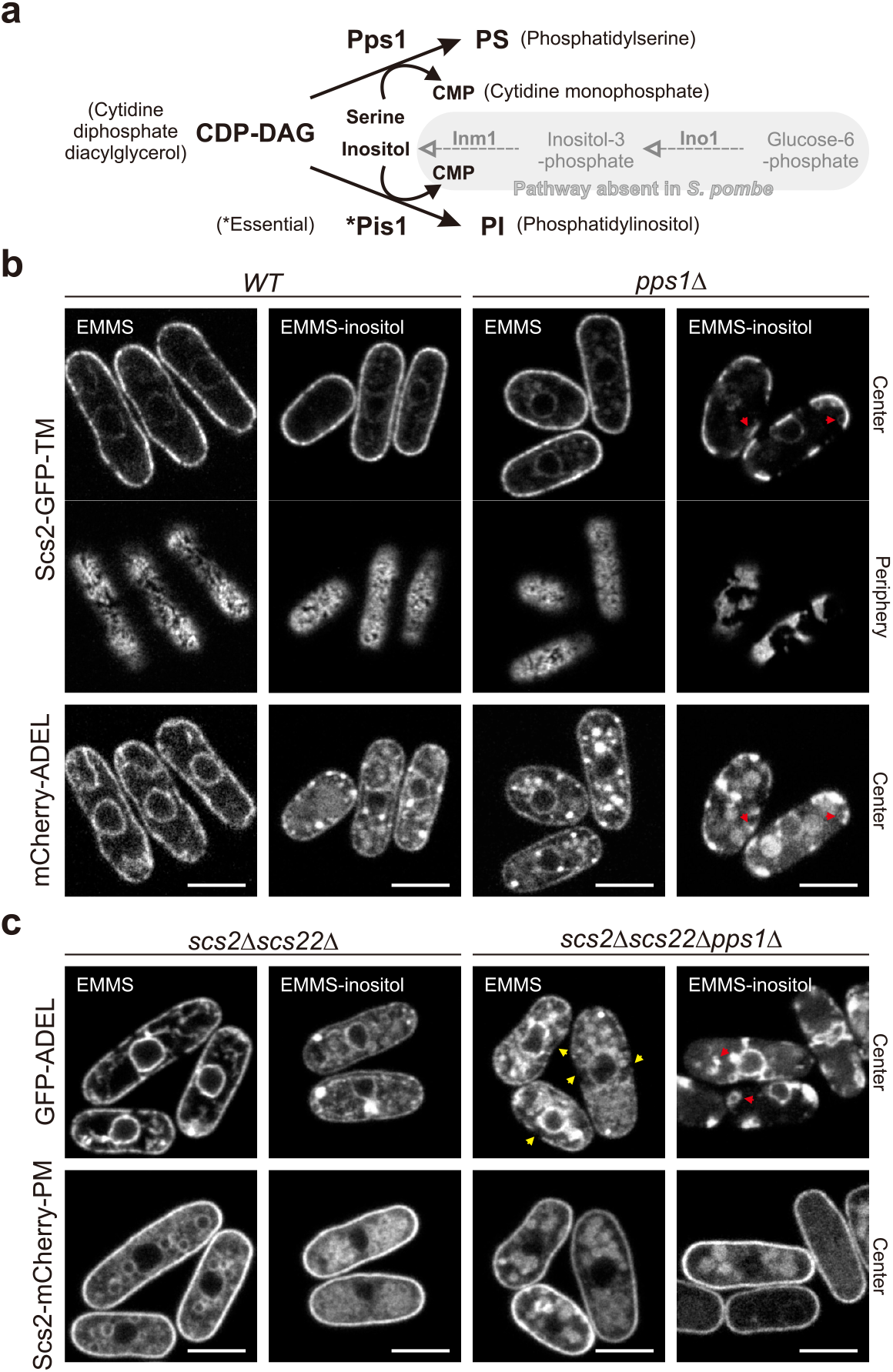
VAP-mediated ER-PM contact formation is compromised in cells lacking inositol and PS. (**a**) Pathways for PI and PS synthesis in fission yeast. *De novo* inositol biosynthesis pathway in *S. cerevisiae* that absent in *S. pombe* is shown in grey. (**b,c**) Scanning confocal images of indicated cells expressing indicated proteins cultured in the presence or absence of inositol. Red arrowheads designate the cytoplasmic ER compartments. Yellow arrowheads point to the cortical regions lacking ER-PM contacts. EMMS, Edinburgh minimal media supplemented with appropriate amino acids. Scale bars, 5 μm.

In fission yeast, PI synthesis is controlled by the essential gene *pis1* that encodes a CDP-DAG-inositol 3-phosphatidyltransferase ^36^ (Fig. 4a). Unlike budding yeast, *S. pombe* is a natural inositol auxotroph lacking the *de novo* synthesis pathway (Fig. 4a) ^37, 38^, which allows us to limit levels of PI and/or PIPs via inositol starvation. Although no method has been established to visually indicate PI abundance in yeast, a gradual decline in cellular PI level was observed after inositol starvation ^39^. Intriguingly, considerably large PM regions were deprived of the ER in inositol-starved *pps1Δ*, but not in such starved *WT* cells (Fig. 4b; Supplementary Movie 1). Both PM levels of PI_4_P and PI_4,5_P_2_ were significantly lowered after inositol starvation (Supplementary Fig. 4). We think that loss of ER-PM contacts specifically seen in inositol-starved *pps1Δ* cells was not due to cell death-related ER mis-organization which also occurred in *WT* after inositol starvation (Fig. 4b). It is rather a consequence of lacking major anionic PLs – the PM counterparts of VAP-mediated ER-PM tethering machinery. Indeed, despite the total amount of the ER may reduce in inositol-starved *pps1Δ* cells, free ER compartments were observed in the cytoplasm (arrowed in Fig. 4b), suggesting that ER-PM contact formation is less favored.

As PI_4_P and PI_4,5_P_2_ are generally scarce in the ER membrane, we further envisaged that ER-PM contacts kept by PM-anchored Scs2 in *scs2Δscs22Δ* backgrounds would be more vulnerable in a shortage of major abundant anionic PLs - PI and PS. Indeed, large ER-free PM regions were readily seen in *scs2Δscs22Δpps1Δ* cells highly expressing Scs2-PM, while these cells mostly lost ER-PM contacts after grown in inositol-free medium, resulting in similarly clustered ER compartments in the cytoplasm (Fig. 4c).

Therefore, we conclude that major anionic PLs, including PI, PS, PI_4_P and PI_4,5_P_2_, collectively act as the essential PM binding partners underpinning VAP-mediated ER-PM tethering in fission yeast.

### VAP-mediated ER-PM contact formation is responsive to cytosolic pH

It has been shown that charge-based interactions of several PH domains with PI_4_P and PI_4,5_P_2_ are pH-sensitive and abolished at low pH (e.g., pH 6 and pH 6.8 for their *in vitro* binding of PH_OSBP_ respectively) ^40^. Our *in vitro* liposome binding results also showed similarly reduced affinities of Scs2N to both PI and PS at a lower pH (pH 5), suggestive of their possible binding through electrostatic attractions (Supplementary Fig. 5a).

We adopted two different approaches to decrease cytosolic pH (pH_c_): 1) inactivation of proton-transporting ATPases by energy depletion (ED) (i.e, by inhibiting both glycolysis and mitochondrial electron transport); 2) treatment with semi-permeable sorbic acid (SA) ^41, 42^. In both cases, pH_c_ in *WT* dropped below 5.5 within two hours indicated by the genetically encoded pH sensor sfpHluorin ^43^ (Supplementary Fig. 5b). Interestingly, both PI_4_P and PI_4,5_P_2_ markers lost the cortical binding after two-hour treatments, while the PS indicator left the cortex only in cells treated with ED but not SA (Fig. 5a). It should be noted that PI_4_P and PI_4,5_P_2_ markers no longer reflect corresponding lipid levels at low pH_c_ where charge-based lipid-marker interactions are compromised ^40^. On the other hand, lipid markers that bind via non-charge-based interactions, e.g., sterols and diacylglycerol (DAG), are possibly pH-insensitive and indeed remained membrane-associated after cytoplasmic acidification (Supplementary Fig. 5c). Localizational difference of the PS marker after treatments should be unrelated to low pH_c_, which also implies a specific role of energy metabolic pathways in maintaining the PM level of PS.

**Figure 5.**
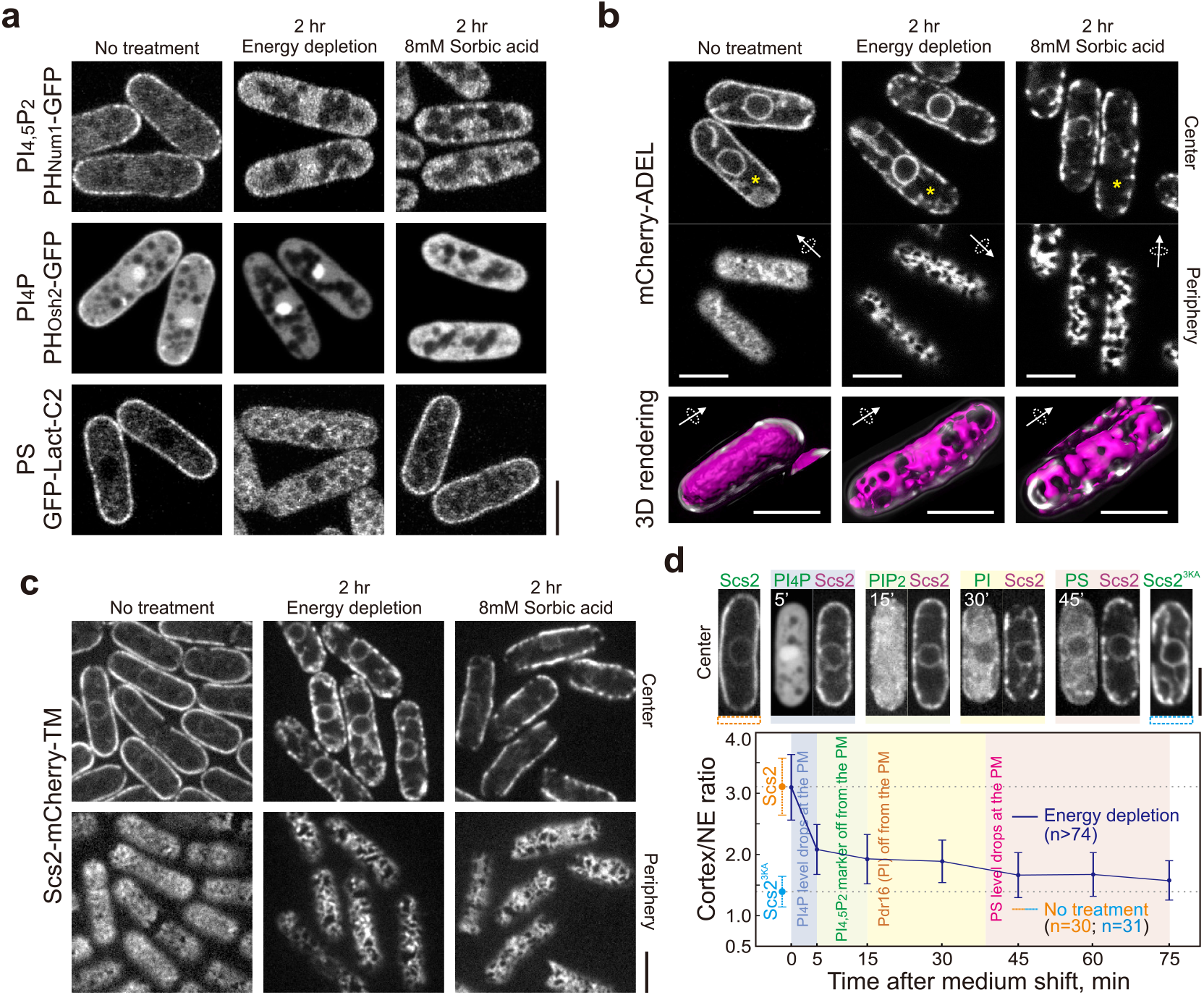
VAP-mediated ER-PM contact formation is compromised in cells with low cytosolic pH. (**a**) Scanning confocal images of *WT* cells expressing various phospholipid markers after indicated treatments. Shown are central focal planes. (**b**) Scanning confocal images of *WT* cells expressing mCherry-ADEL after various treatments. The corresponding 3D-rendered cortical snapshots of asterisked cells are shown in the bottom. (**c**) Spinning disk confocal images of *WT* cells expressing Scs2-GFP-TM after indicated treatments. (**d**) Reduction in cortex/NE ratios of Scs2-mCherry-TM after energy depletion in *WT* cells. Shown are representative central focal plane spinning disk confocal images of indicated cells with or without the treatment. Time, relative to buffer shift. Cortex/NE ratios of Scs2-GFP-TM and Scs2^3KA^-GFP-TM from non-treated cells are included as controls. Time windows for the initiation of each indicated events are highlighted with corresponding colors. Error bars represent 2×SD. n, cell number. Scale bars, 5 μm.

Importantly, ER-PM contacts were significantly reduced in these *WT* cells after both treatments (Fig. 5b; Supplementary Movie 2). Notably, PM affinity of Scs2 was greatly reduced in energy-depleted cells to almost Scs2^3KA^ level within two hours, manifested by decreased cortex-to-NE ratios (Fig. 5c,d). Such affinity reduction occurred progressively as PI_4_P, PI_4,5_P_2_ and PS markers sequentially lost the PM localization (Fig. 5d). As PI_4_P marker left the cortex immediately upon cell exposure to ED buffer before possible pH_c_ decline, we favored the idea that these cells lost the PM pool of PI_4_P at the early stage with unknown mechanisms. PM affinity of Scs2 did not obviously reduce in SA-treated cells (Fig. 5c), which was not simply due to the presence of PS in the PM (Fig. 5a), as *pps1Δ* lacking PS also exhibited similarly after SA treatment (Supplementary Fig. 5d). We guessed that either PM PI levels or PI-Scs2 interactions could be affected differently in ED- and SA-treated cells. Somewhat as indirect evidence, we found that Pdr16, a presumed PI transporter ^44^, lost the PM binding only in cells treated with ED but not SA (Supplementary Fig. 5c). In fact, it left the cortex prior to the PS marker in these cells (Fig. 5d), which may reflect reduced PI availability in the PM after energy depletion. We confirmed that fission yeast Pdr16 bound PI *in vitro* (Supplementary Fig. 5e). Concordantly, PM localization of Pdr16 was barely visible in inositol-starved *pps1Δ* cells which should have lost most PI (Supplementary Fig. 5f).

Collectively, these results again demonstrate our view that the cortical binding of Scs2 indeed depends on interactions with major anionic PLs, especially PI and PS. It further suggests that lipid-VAP interaction-based ER-PM contacts are responsive to cytosolic pH.

### Scs2 modulates Pma1 via FFAT-mediated interaction for pH homeostasis

Given that the major pH_c_ regulator Pma1 was among the top hits of Scs2-interactors (Fig. 2b), we wondered if Scs2 plays a role in pH_c_ homeostasis. Interestingly, we identified a highly conserved FFAT-like motif within the P domain of Pma1 across fungi (Fig. 6a). Either the removal or genetic alterations of this presumed Scs2-interacting motif engendered inviable spores in *S. pombe* (Fig. 6a), indicative of its necessity for Pma1 function. We confirmed Scs2-Pma1 interaction *in vivo* using split-YFP-based bimolecular fluorescence complementation assay ^45^ (BiFC; Fig. 6b). Scs2^3KA^ lacking the functional MSP domain that is known FFAT motif-associated ^22^, did exhibit much weaker binding (if any) to Pma1 indicated from both Co-IP and BiFC experiments (Supplementary Fig. 6a,b).

**Figure 6.**
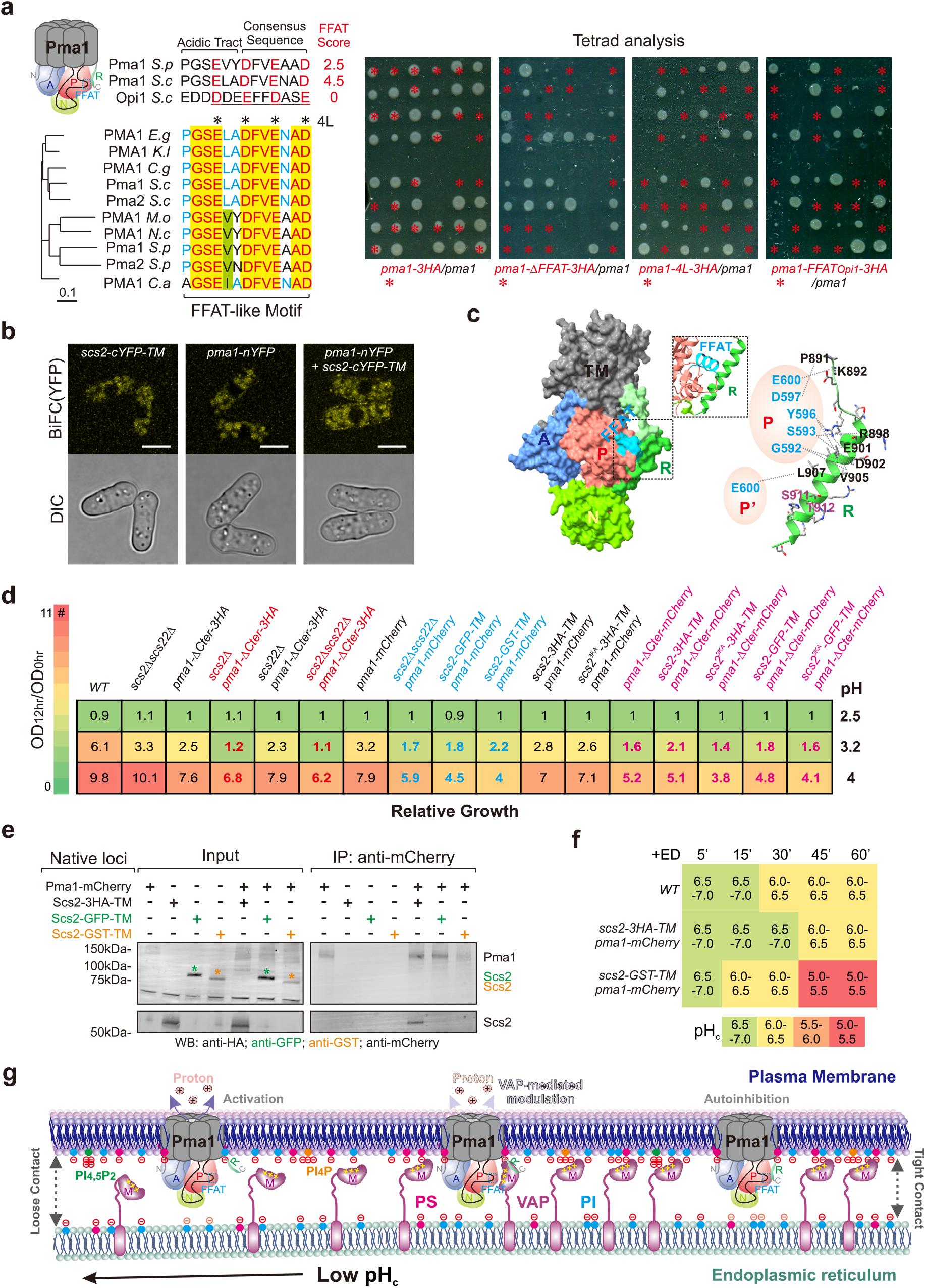
Scs2 modulates Pma1 activity via FFAT-like motif-mediated interaction for pH_c_ homeostasis. (**a**) Predicted FFAT-like motif locating within the P domain of Pma1 is highly conserved in Fungi and essential for Pma1function in fission yeast. *S.p, S. pombe; S.c, S. cerevisiae; E.g, E. gossypii; K.l, K. lactis; C.g, C. glabrata; M.o, M. oryzae; N.c, N. crassa; C.a, C. albicans*. The underlined Opi1 sequence is the part replacing the corresponding Pma1 segment in *pma1-FFAT_Opi1_* mutant. Black asterisks mark mutated residues and red asterisks point to colonies or inviable spores carrying indicated pma1 mutants. (**b**) Central focal plane scanning confocal images of cells expressing indicated proteins. Scale bars, 5 μm. (**c**) Predicted structure of fission yeast Pma1 monomer (108E-918D). TM (transmembrane), A (actuator), N (nucleotide-binding), P (phosphorylation), and R (regulatory) domains are indicated. Contact between R domain and FFAT-like motif is illustrated in the frame. Predicted intramolecular (R-P) and intermolecular (R-P’of the adjacent monomer) interactions during R domain-mediated autoinhibition are shown with dotted lines. Involved residues are conserved between *S. pombe* and *N. crassa.* S911 and T912 are conserved phosphorylation sites for autoinhibitory regulation. (**d**) Growth assays of indicated cells in medium with different pH. Relative growth (OD_12hr_/OD_0hr_) over 12 hours is indicated by both number and color. pH-sensitive strains were highlighted in colors. (**e**) Co-IP of indicated proteins expressed from native loci. Samples were probed with indicated antibodies. (**f**) pH_c_ of sfpHluorin-expressing cells with indicated backgrounds at different time points after energy depletion (ED). The times are relative to ED buffer addition. pH_c_ range is indicated by both number and color. For each cell type, n>170 cells were used for measurement. (**g**) A model for mechanisms of VAP-mediated ER-PM contact formation and yeast VAP function in pH_c_ homeostasis via Pma1 modulation. M, MSP domain.

Recently resolved Pma1 structures in both *S. cerevisiae* and *N. crassa* have pinpointed autoinhibitory regulation of Pma1 through intra- or inter-molecular interactions between P and R domains ^15, 16^. Remarkably, the FFAT-like motif is right at this P-R interface (Fig. 6c), implicating Scs2 in Pma1 regulation. In fact, *scs2Δscs22Δ* cells already showed compromised growth in highly acidic media, similarly to *pma1-ΔCter* cells that lacked the C-terminal R domain (Fig. 6d). Moreover, Scs2 appeared to play a specific role in surviving extreme acid environments, as *pma1-ΔCter* cells became more sensitive to low pH when they further lost Scs2 (Fig. 6d).

Surprisingly, *WT* cells expressing both Scs2-GFP-TM and Pma1-mCherry were sensitive to highly acidic media, similarly to that of Pma1-mCherry-expressing *scs2Δscs22Δ* cells (Fig. 6d). These data suggest that pH-related growth defects associated with *scs2Δ* were not simply due to ER-PM contact loss. Intriguingly, Scs2-Pma1 interaction was totally abolished in these Pma1-mCherry-expressing *WT* cells where Scs2 was GFP-tagged (Fig. 6e). In contrast, such growth defects were not seen in *WT* cells when Scs2 was tagged with 3HA and Scs2-Pma1 interaction was retained (Fig. 6d,e). Thus, Scs2-Pma1 interaction is important for cell survival in acid environments. Besides, Pma1-mCherry localization was unaffected in all these backgrounds regardless of functional VAPs or Scs2-Pma1 interaction (Supplementary Fig. 6c), indicating that Scs2-mediated modulation is non-essential for wild-type Pma1 trafficking to the PM. On the other hand, Pma1-ΔCter-mCherry localized normally to the PM in the presence of Scs2-Pma1 interaction. However, it became defective in the ER exit without this interaction (Supplementary Fig. 6d), implying that Scs2 binding might also aid proper Pma1 assembly or maturation.

We then imagined that Scs2-mediated modulation on Pma1 activity could be crucial for pH_c_ homeostasis. To directly test this idea, we took advantage of brief ED treatment which stops glucose-dependent elimination of Pma1 autoinhibition ^17^ and meanwhile prohibits the production of ATP that powers Pma1 to maintain pH_c_ ^46^. We traced pH_c_ decline in ED-treated cells over time. As expected, unlike in *WT*, pH_c_ dropped rapidly below six within one hour in cells lacking either or both of functional Scs2 and Pma1 R domain (Supplementary Fig. 6e). Of note, in these ED-treated *WT* cells, Scs2-Pma1 interaction remained after mild cytoplasmic acidification (Supplementary Fig. 6f). We then wondered if Scs2-Pma1 interaction is required for sustaining pH_c_ of cells facing low glucose and ATP with brief ED treatment. To this end, we firstly generated *WT* cells expressing both Scs2-GST-TM and Pma1-mCherry where Scs2-Pma1 interaction was confirmed similarly absent as in *WT* background where Scs2 was GFP-tagged (Fig. 6e). These cells were also similarly sensitive to acidic media (Fig. 6d). Indeed, pH_c_ dropped rapidly below six in these cells within one-hour ED treatment, unlike *WT* backgrounds where Scs2-Pma1 interaction was intact (Fig. 6f). Collectively, our results suggest that FFAT-mediated interaction enables Scs2 to functionally modulate Pma1 for robust pH_c_ homeostasis, likely by hijacking Pma1 autoinhibition via a competitive binding to its P domain.

Altogether, we conclude that conserved interactions between VAPs and major anionic phospholipids underlie VAP-mediated ER-PM contact formation and hence determine the pH-sensitive nature of ER-PM contacts in fission yeast. We also hypothesize that Scs2 maintains cytosolic pH through its modulatory interaction with Pma1 (Fig. 6g). All such interactions necessitate the conserved MSP domain.

## Discussion

Despite lacking concrete evidence, VAP-protein interactions taken place at various MCSs, especially the ones bridged by FFAT or related motifs, have been widely regarded as primary ‘molecular pillars’ underlying VAP-mediated membrane tethering ^1, 2^. Here, we provided both *in vitro* and *in vivo* evidence to postulate an alternative viewpoint that VAP-phospholipid (e.g., PI and PS) interactions are the major molecular forces at least for ER-PM coupling in fission yeast. Conceivably, major anionic PLs of the PM (comprised of ∼ 20% PI, 10% PS and 1% PI_4_P + PI_4,5_P_2_ in yeast ^31, 46^) are in much higher abundance than VAPs, which makes the extent of ER-PM contacts dependent on VAP dosage. We believe that such tethering mechanism should not be merely limited to *S. pombe* or to ER-PM contacts, as binding of anionic PLs appears to be conserved in VAP family and other organelles may also contain adequate levels of anionic PLs in cytosolic leaflets ^30, 47^. Of note, budding yeast Scs2 was previously shown to bind phosphoinositides and PS *in vitro* ^48^, however the study did not link such phospholipid affinities to the membrane tethering ability of Scs2. Moreover, much lower PI content in the mammalian PM as compared to that of yeasts ^30, 49, 50^ and variation in PS-binding ability of VAPs may partially explain different scales of ER-PM association in these cell types. On the other hand, distribution of these anionic PLs among organelles could in turn define preferential sorts of VAP-mediated ER contacts for a given cell type under various conditions.

VAPs facilitate recruitment of several ORP/Osh proteins and subsequently their non-vesicular sterol/PI_4_P transfer at ER-PM/Golgi contact sites to allow ER-resident Sac1-mediated PI_4_P metabolism and hence sustain PI_4_P levels in respective membranes ^9, 51^. Of interest, our data suggest that the ER localization of VAPs is no longer required for this PI_4_P metabolism in fission yeast when VAP-mediated ER-PM contacts are intact. Thus, VAPs likely have no impact on directionality of such lipid transfer at MCSs. It has been speculated that ORP/Osh-dependent sterol/PI_4_P countertransport and Sec14 family-mediated PI/PC exchange may be functionally coupled at ER-PM contacts ^52^. Here we showed that the cortical localization of Sec14-like Csr102 is fully dependent on VAPs, which somewhat substantiates the possibility of such interconnected lipid exchange cycles via VAPs, although exact functions of Csr102 still await further investigation. Additionally, VAPs’ affinity to a wide range of anionic PLs could further make them an ideal regulatory nexus to coordinate multiple lipid exchange reactions at MCSs. It is thus attractive to speculate that ALS-associated VAPB^P56S^ with reduced affinity to anionic PLs may fail such coordination hence promoting disease progression.

Ionized status of anionic PLs is pH-dependent, thus their electrostatic interactions with proteins are pH-sensitive and mainly prevail at pHs above respective pKa. For instance, as pKa values for the 4-phosphate moiety of PI_4_P (∼6.3) and PI_4,5_P_2_ (∼6.7) ^53^ fall within the physiological range, their interactions with proteins are vulnerable to cytosolic acidification ^40^. Similarly, we saw a sequential loss of PM binding for PI_4,5_P_2_ and PI_4_P markers in SA-treated cells as pH_c_ decreased (data not shown). However, ionization of PI (pKa ∼2.5) and PS (pKa ∼5.5) is less likely affected by physiological pH_c_ oscillation ^54^. Indeed, protein binding of PI and PS remained when pH_c_ dropped below 5.5 in SA-treated cells. As discussed, we think that available PM pools of PI and PS are specifically reduced after energy depletion. Conceivably, ATP consumption that drives lipid conversion and circulation is likely required to maintain PI and PS levels in the PM.

Glucose/energy sensing is intimately linked to Pma1 regulation. It is widely accepted that the C-terminal R domain is autoinhibitory at normal pH_c_ by immobilizing other cytosolic domains via a clamp-like interaction with the P domain ^15, 16, 55–58^. By releasing the cytosolic domains, phosphorylation of the R domain upon glucose sensing rapidly activates Pma1 ^17, 59^. Our study suggests a second regulatory module in interfering Pma1 autoinhibition via an FFAT-VAP-mediated competitive binding to the P domain. This module could act to prolong Pma1 activity in cells facing low glucose to avoid rapid cytoplasmic acidification before ATP exhaustion. Ultrastructural insights will help to address if such interactions are constantly stimulative or versatile dependently on conditions. We conceive that such modulatory VAP-Pma1 interaction mainly works *in trans* across ER-PM contacts. However, a certain portion of their *cis* interactions may facilitate Pma1 assembly in the ER and its proper packaging for ER exit, especially when it lacks the R domain. It remains to be seen if VAPs play such a general role in protein trafficking, considering they interact widely with vesicle trafficking machinery.

Of note, PS has been shown enriched in the PM regions surrounding transmembrane proteins, including that of Pma1 ^31^. A recent study also found that several TMs of Pma1 exhibted a preferred binding with PS in the inner PM leaflet ^16^. It is thus likely that PS binding of Scs2 may facilitate its access to the vicinity of Pma1 for the modulatory interaction. Another enthralling idea is that pH_c_-sensory interactions of Scs2 with PI_4_P and PI_4,5_P_2_ might autonomously signal its favored binding towards Pma1 when pH_c_ slightly drops. We therefore hypothesize that VAPs balance pH_c_ by gearing their preferential interactions between Pma1 and pH-biosensory PLs at ER-PM contacts. Since both interactions appear conserved, this strategy could be widely shared by other organisms.

## Methods

### Experimental Model and Subject Details

All fission yeast strains used in this study are listed in Supplementary Table 1. Strains were constructed from laboratory stocks using standard methods. The artificial ER-PM tether (TM-GFP/mCherry-CSS_Ist2_) and the control construct (TM-GFP/mCherry) established previously in ^5^ were expressed from *leu1* locus under Thiamine-repressible *nmt1* promoter. The PI_4_P marker PH_Osh2_-mCherry was designed as PH_Osh2_-GFP previously reported in ^5^ and was constitutively expressed from *ura4* locus under *rtn1* promoter. The PS marker GFP-Lact-C2^35^ was expressed from *ade6* locus under *act1* promoter, the sterol marker GFP-D4H ^60^ was expressed from *leu1* locus under *pil1* promoter, and the DAG maker C1δ-GFP amplified from rat PKCδ C1-containing plasmid (Addgene #21216) ^61^ was expressed from *ura4* locus under *scs2* promoter. The pH sensor superfolder pHluorin (sfpHluorin) amplified from p426MET25-sfpHluorin (Addgene #115697) ^43^ were expressed from *leu1* locus under *pil1* promoter.

The strain where *scs2* and *scs22* swapped their genomic loci shown in Supplementary Fig. 1a was made by replacing the entire open reading frame of one gene with the other (inclusive of introns for both genes). Scs2-3HA-TM and Scs22-3HA-TM were generated by inserting three tandem HA epitopes between S360 and P361 of Scs2 or between I296 and H297 of Scs22 that expressed from their respective native locus under their own promoters, in the same way as previously described ^5^. Scs2-mCherry-PM or Scs2^3KA^-mCherry-PM was generated by inserting mCherry after S360 and by replacing TM region with the PM targeting motif (“KKKKKKSKTKCVIM”) from human K*-*Ras4B ^29^. It was expressed from *leu1* locus under either *nmt1* or *pil1* promoter. VAPB-GFP-TM or VAPB^P56S^-GFP-TM was generated by inserting GFP between P213 and R214 of VAPB or VAPB^P56S^ that expressed from *leu1* locus under *pil1* promoter. GFP-tagged Scs2 and Scs22 were expressed from respective native locus unless otherwise stated. In *scs2Δscs22Δ* background, as annotated in Fig. 1-2 and Supplementary Fig. 1-3, GFP-tagged Scs2 and Scs22 were expressed from *leu1* locus under *scs2* or *pil1* promoter, and GFP-tagged mutants and swop variants (i.e., Scs2^3KA^ or Scs2^K36/38/43A^, Scs2^2TA^ or Scs2^T39/40A^, Scs22^3KA^ or Scs22^K36/38/43A^, MSP_scs2(1-125)_-LK-TM_scs22(122-319)_, and MSP_scs22(1-121)_-LK-TM_scs2(126-383)_) were expressed from *leu1* locus under *pil1* promoter. C-terminally GFP tagged N-terminus (lacking only the TM domain) of either Scs2 (i.e., Scs2N or Scs2N^1–360^) or Scs22 (i.e., Scs22N or Scs22N^1–296^) used for proteomics analyses in Fig. 2 and Supplementary Fig. 2 was expressed from respective native locus. C-terminally GFP tagged N-terminus of VAP variants (namely Scs2N, Scs22N, Scs2N^3KA^, Scs2N^2TA^, Scs22N^3KA^, VAPBN^1–213^ and VAPBN^1–213^(P56S)) used for liposome analyses in Fig. 3 and Supplementary Fig. 3,5 were expressed from *leu1* locus under *pil1* promoter in *scs2Δscs22Δ* cells. In Supplementary Fig. 3a, GFP-tagged Scs2 mutants (i.e., Scs2^3KA^, Scs2^TA^) were expressed from the native locus. Scs2-cYFP-TM were generated by inserting cYFP between S360 and P361 of Scs2 that expressed from its native locus. N-terminally nYFP tagged Scs2 was expressed from *leu1* locus under *scs2* promoter.

C-terminally tagged Csr102 or Pdr16 were expressed from their respective native locus. C-terminally 3HA, mCherry, nYFP and cYFP tagged Pma1 mutants were all expressed from the native locus. Mutations located in the FFAT-like motif of Pma1 resulted in spontaneous diploidization. Pma1-ΔFFAT was made by a deletion of 13 amino acids from 591P to 603D; Pma1-4L was made by substituting all 594E, 597D, 600E and 603D with leucine; Pma1-FFAT_Opi1_ was constructed by replacing 10 amino acids from 594E to 603D by “DDEEFFDASE” from budding yeast Opi1. Pma1^ΔCter^ was generated by deleting its last 23 amino acids (897Q-919A).

For all experiments unless otherwise stated, *S. pombe* cells were grown exponentially at 24 °C in YES (yeast extract with supplements) or EMMS (Edinburgh minimal media supplemented with appropriate amino acids) before specific treatments or imaging. *pps1Δ* cells were freshly grown on YES medium plate supplemented with 0.4 mM ethanolamine (Merck, 398136-25ML) before liquid culture. Expression of artificial constructs under *nmt1* promoter was induced in EMMS for at least 20 hours. EMMS with 5 mg/mL thiamine (Thi) (Sigma-Aldrich, T1270-25G) was used to suppress the construct expression. For inositol starvation experiments, cells were cultured in EMMS without inositol for at least 21 hours.

### Microscopy

Scanning confocal microscopy was performed on an Olympus FLUOVIEW FV3000 (Olympus Corporation, Japan) equipped with a U Plan Super Apochromat 100X, 1.4 NA Oil immersion objective lens, a 488 nm (for GFP excitation), a 514 nm (for YFP excitation) and a 561 nm (for mCherry excitation) solid state laser, with high sensitivity-spectral detectors. Typically, we acquired image stacks that consisted of 21 z-sections with 0.24 µm spacing or 9 z-sections with 0.5 µm spacing (55.2 nm per pixel).

We also employed Nikon TiE system (CFI Plan Apochromat VC 100X, 1.4 N.A. objective) equipped with Yokogawa CSU-X1-A1 spinning disk unit, the Photometrics CoolSNAP HQ2 (129 nm per pixel with 2 × 2 binning) or Prime 95B™ Scientific CMOS camera (110 nm per pixel) and a DPSS 405 nm 100 mW, a DPSS 491 nm 100 mW and DPSS 561 nm 50 mW laser illumination under the control of MetaMorph Premier Ver. 7.7.5 to collect single or multiple z-plane images or to perform fluorescence recovery after photobleaching (FRAP) experiments. All FRAP experiments were done in the same microscopy setting. Typically, a 10 × 10 pixels circular region at the cell cortex (top plane) was bleached with a sequence of 200 high-intensity laser iterations and then subsequent images were taken every 0.5 second for 25 seconds (Supplementary Fig. 1b,c; Pdr16 and Csr102 in Supplementary Fig. 2i) or 1 minute for 5 minutes (Pma1 and Pil1 in Supplementary Fig. 2i).

Unless otherwise stated, imaging was performed on *S. pombe* cells placed in sealed growth chambers containing 2% agarose YES medium or indicated growth medium. Only interphase cells were used in all image analysis in this paper.

### Proteomics for VAPs, protein level estimation and Co-immunoprecipitation (Co-IP)

Cells expressing Scs2N^1–360^–GFP, Scs2-GFP-TM, Scs22N^1–296^–GFP, Scs22-GFP-TM or GFP control were grown to log phase, harvested with freshly prepared buffer A (50 mM Tris-HCl, 150 mM NaCl, 2 mM EDTA, 1% Nonidet P (NP)-40, 50 mM NaF, 0.1 mM Na_3_VO_4_, 1 mM PMSF, 1.5 mM Benzamidine-HCl, and Roche protease inhibitor cocktail). Cells were mixed with glass beads and homogenized in a Mini Bead Beater (Biospec, OK, USA) at 4 °C and total cell lysates were harvested after removing cell debris. Lysates were adjusted to the same total protein concentration using buffer B (10 mM Tris-HCl, 150 mM NaCl, 0.5 mM EDTA, 50 mM NaF, 0.1 mM Na_3_VO_4_, pH 7.4, added with protease inhibitors) before incubation with GFP-Trap beads (Chromotek, Munich, Germany) for one hour at 4 °C. Beads were washed with buffer B and resuspended in SDS loading buffer. Co-immunoprecipitated proteins were visualized with Coomassie staining reagent (Bio-Rad), and excised gel segments or total Co-immunoprecipitated proteins in solution were sent for LC-MS/MS analysis. All mass spectrometry was performed by Protein and Proteomics Center (PPC) in the National University of Singapore. We preformed three replicates with Scs2 and Scs22, and two replicates with Scs2N and Scs22N. GFP control was used as the reference for gel excision and its MS data were used for further excluding non-specific interactors in interactome analysis. Data and quantification shown in Fig. 2 and Supplementary Fig. 2 were from one of three replicated IP-MS experiments. Interactors were defined using a minimum peptide count of four.

For protein level estimation in Fig. 1b,3e and Supplementary Fig. 1d, total cell lysates were harvested as described above, and then subjected SDS-PAGE and western blotting probed by rabbit anti-GFP (Abcam, MA, USA) and mouse anti-α-tubulin antibodies (TAT-1, a gift from K. Gull, University of Oxford, UK). For detection, IRDye800 conjugated anti-mouse and IRDye680 conjugated anti-rabbit antibodies were used and subsequent quantification analyses were done on the Odyssey infrared imaging system (LI-COR, NE, USA) and ImageJ 1.48v software package (NIH, MD, USA).

For co-immunoprecipitation, glass bead-homogenized cells or cryo-ground (only for Co-IP experiments with Pil1-mGFP) were suspended in 300 µL of buffer A at 4 °C and total cell lysates were harvested after removing cell debris. We used low salt buffers where NaCl concentration was 50 mM for Co-IP experiments with Pil1-mGFP and Pma1-mCherry. Typically, lysates were adjusted to the same total protein concentration, diluted with buffer C (the same recipe as buffer A but without NP-40) and precleared for non-specific binding by incubation with Dynabeads^TM^ Protein G (Invitrogen, CA, USA) for one hour at 4 °C. The pre-cleared lysate was then subjected to incubation with rabbit anti-GFP, rabbit anti-mCherry (Abcam, MA, USA), or mouse anti-HA (Roche, Basel, Switzerland) antibody for one hour at 4 °C and then with Dynabeads^TM^ Protein G for 45 minutes at 4 °C. Beads were washed five times with buffer C and additionally one time with 1% NP-40-containing buffer A. We also repeated experiments shown in Fig. 6e and Supplementary Fig. 6a,f with ChromoTek RFP-Trap magnetic beads (Proteintech, IL, USA). Samples were resuspended in SDS-loading buffer and subjected SDS-PAGE and western blotting probed by rabbit anti-GFP, rabbit anti-mCherry or mouse anti-HA antibodies.

### Recombinant protein purification

Scs2N-6His-expressing plasmid-containing BL21 (DE3) cells were cultured in Lysogenia Broth (LB) medium at 37°C to an OD_600_ of 0.7-1.0. Protein expression was then induced by 0.5 mM IPTG for 5-6 hours or overnight at 24°C. Cells were lysed by sonication in buffer D (50 mM NaH_2_PO_4_ (pH 8), 500 mM NaCl, 40 mM Imidazole, 2 mM PMSF, 0.001 g/mL lysozyme) and debris were removed by centrifugation at 20,000 g for 15 mins at 4°C. Scs2N-6His was purified using the standard protocol with the Thermo Scientific™ HisPur™ Ni-NTA spin column.

### Liposome preparation

Lipids were purchased from either Sigma-Aldrich: L-α-phosphatidylcholine (PC, egg yolk) or Avanti Polar Lipids: L-α-phosphatidylinositol (PI, soy); L-α-phosphatidylserine (PS, brain); L-α-phosphatidylinositol-4-phosphate (PI_4_P, brain); L-α-phosphatidylinositol-4,5-biphosphate (PI_4,5_P_2_, brain); and 1,2-dioleoyl-sn-glycero-3-phosphoethanolamine-N-(lissamine rhodamine B sulfonyl) (Rhod-PE). Lipids were dissolved and mixed in chloroform and dried using SpeedVac vacuum concentrator (Eppendorf, HH, DE) for 24 hours.

For co-sedimentation assays, respective lipid films (i.e., 100% PC (mol/mol); 75% PC and 25% PI; 20% PC and 80% PI; 75% PC and 25% PS; 20% PC and 80% PS; 73% PC, 25% PI and 2% PI_4_P; 73% PC, 25% PS and 2% PI_4_P) were hydrated with buffer E (20 mM HEPES, 150 mM NaCl, 0.75 M sucrose, pH 7) and sonicated for 20 mins at 65°C. Liposomes were passed through polycarbonate membrane with a pore size of 100 nm using the MiniExtruder (Avanti Polar Lipids, AL, USA) at 65°C. Liposomes were washed with buffer F (20 mM HEPES, 150 mM NaCl, pH 7) and concentrated by centrifugation at 32,000 g for 30 mins at 4°C.

For binding assays, respective 1% Rhod-PE containing lipid films (i.e., 99% PC; 74% PC and 25% PI; 74% PC and 25% PS; 72% PC, 25% PI and 2% PI_4_P; 72% PC, 25% PI and 2% PI_4,5_P_2_; 72% PC, 25% PS and 2% PI_4_P; 72% PC, 25% PS and 2% PI_4,5_P_2_) were hydrated with buffer F or G (the same recipe as buffer F but with a pH of 5) and sonicated for 20 mins at 65°C. Aggregates were removed by centrifugation at 21,100 g for 30 mins at 4°C. Liposome concentration was normalized by the rhodamine intensity of 2.5% NP-40-lysed liposomes using the Spark multimode microplate reader (Tecan, Männedorf, Switzerland).

### Liposome co-sedimentation assay

Purified Scs2N-6His was incubated with respective liposomes for 30 mins at 4°C with gentle agitation. The mixture was pelleted by centrifugation at 32,000 g for 30 mins at 4°C. Both the supernatant and pellet (after washing with buffer F) were resuspended with SDS-loading buffer for SDS-PAGE and western blotting probed by mouse anti-His antibody (Abcam, MA, USA).

### Liposome binding assay

Normalized soluble lysates of *WT* cells expressing Pdr16-GFP or *scs2Δscs22Δ* cells expressing either Scs2N-GFP, Scs2N^3KA^-GFP, Scs2N^2TA^-GFP, Scs22N-GFP, Scs22N^3KA^-GFP, VAPBN-GFP or VAPBN^P56S-^GFP were freshly prepared using buffer C and incubated with rabbit anti-GFP antibody for one hour and then with Dynabeads^TM^ Protein G for one hour at 4°C with gentle agitation. Dynabeads conjugated with saturated protein of interest were washed with buffer C before incubation with normalized Rhod-PE-containing liposomes for one hour at 4°C. Soluble *WT* or *scs2Δscs22Δ* cell lysates were used as the negative control respectively. liposomes were prepared with buffer G for assays preformed under pH 5. Dynabeads were harshly washed with buffer F or G and imaged by confocal microscope on 96-well plates.

### Energy depletion and sorbic acid treatment

Freshly washed cells were incubated on rotary shaker with 1 mL 8mM sorbic acid/EMM or energy depletion buffer (EMM without glucose, 20 mM 2-geoxyglucose (2-DG), 10 µM antimycin A) before imaging. For confocal imaging, cells were placed on an agarose pad prepared from respective media.

### Cytosolic pH measurement

Cytosolic pH of sfpHluorin-expressing cells after various treatments was indicated from fluorescence intensities obtained by confocal microscopy using DAPI/GFP (Excitation: 405 nm; Emission: 488 nm) and GFP/GFP (Excitation and emission: 488 nm) filter sets as previously described ^41^. DAPI/GFP to GFP/GFP intensity (simplified as 405/488) ratios were used to designate the cytosolic pH against the standard calibration curve. To generate the calibration curve, sfpHluorin-expressing cells were treated with calibration buffers (50 mM MES, 50 mM HEPES, 50 mM KCl, 50 mM NaCl, 0.2 M ammonium acetate, 75 mM monensin, 10 mM nigericin, 10 mM 2-DG and 10 mM NaN_3_) of various pH (5.15, 5.51, 6.03. 6.65, 7.17, and 7.73 in Supplementary Fig. 5b; 5.0, 5.5, 6.0. 6.5, 7.0, and 7.5 in Fig. 6f and Supplementary Fig. 6e). As previously described ^62^, calibration buffers allowed rapid equilibration of the intracellular and extracellular pH. The standard curve was made by plotting 405/488 ratios of these cells under each pH condition against respective pH.

### pH sensitivity growth assay

Cells were firstly cultured in YES till exponential growth phase. ∼0.05 OD_595_ of fresh cells were used for each growth assay performed on the 24-well plate: cells were grown in YES of respective pH (2.5, 3.2, and 4) at 30 °C (1.5 mL per well). OD_595_ was measured by the Spark multimode microplate reader every 6 hours for 12 hours.

### Image processing

#### Intensity profile analysis

For intensity profile analysis shown in Fig. 2d,3b, we selected a non-medial region (avoiding the nucleus) using a line of 40-pixel width perpendicular to the long cell axis at central focal plane of a cell and obtained initial intensity the profile of PH_Osh2_-GFP using “Plot Profile” built-in plugin in ImageJ. Intensity corrected by background subtraction was normalized respectively as ÎF = IF/IF_Max_, where IF_Max_ represents the maximum corrected intensity along the line. Normalized intensity and positional information were replotted using MATLAB with a moving average (n=2). Only interphase cells were used for the analysis.

#### 3D rendering

For 3D renderings shown in Fig. 5b and Supplementary Movies 1,2, z-stack images were taken with 0.24 µm spacing on Olympus FLUOVIEW FV3000. Image deconvolution was performed using adaptive point spread function module in Autoquant (Media Cybernetics, Bethesda, MD). 3D rendering was carried out using Imaris software module (Bitplane, South Windsor, CT, USA).

### Quantification and statistical Analysis

#### Measurement of cortex/NE ratio of VAP variants

For each analyzed cell, regions of the cortical ER and the nuclear envelope (NE) were manually outlined according to the localization of either GFP (in Fig. 1c,e and Supplementary Fig. 3a) or mCherry-tagged (in Fig. 5d) VAP variants or the ER luminal marker mCherry-ADEL (in Fig. 1c,e and Supplementary Fig. 3a) at the central focal plane. Mean intensities of respective regions were measured by ImageJ and corrected by background subtraction. The ratio between corrected intensities of corresponding regions was defined as cortex/NE ratio. In particular, cortex/NE ratios of VAP variants shown in Fig. 1c,e and Supplementary Fig. 3a were normalized by dividing corresponding cortex/NE ratios of mCherry-ADEL of the same cell. Only interphase cells were used for the analysis.

#### FRAP analysis

Average fluorescence intensity of the bleached region was corrected for both the background and non-specific bleaching that based on the intensity of a non-bleached cell in the same filed. Corrected intensity for each time point was normalized as ÎF(t) = [IF(t) – IF(0)]/[IF(-Δt) – IF(0)], where IF(-Δt) represents intensity before photobleaching and IF(0) is the first intensity following bleaching. Normalized recovery curves were fitted with a simple single exponential function IF(t)=F(1-e^-kt^) in MATLAB. Recovery half time was calculated as ln(2)/k for Supplementary Fig. 1b,c,2i.

#### Proteome-wide FFAT scoring

A sequence unit of every continuous 13 residues mimicking a FFAT-like motif (i.e., six flank residues preceding seven core amino acids) was used for scoring based on the position weight matrix as previously described ^4, 23^. For each given protein, the FFAT score shown in Fig. 2b,c,6a and Supplementary Fig. 2a,d was the minimal score obtained by scanning over the entire protein sequence. The scoring system was coded in Python and available upon request.

#### Liposome binding analysis

##### Calculation of relative liposome binding

We measured mean GFP and Rhodamine intensities for each analyzed Dynabeads using ImageJ as *F_protein_* and *F_liposome_* respectively. For each liposome type (25% PI or 25% PS), average values of *F_protein_* and *F_liposome_* from all beads incubated with control lysates (i.e. 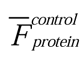 and 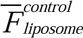) were used for correction, and relative liposome binding of the tested protein for each bead was calculated as (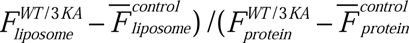) or 0 (if 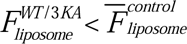) shown with box plots in Fig. 3c and Supplementary Fig. 3e,g. Pseudo-colored images were all with the same contrast for each fluorescent channel. We set the highest rhodamine intensity obtained from the binding assay of the corresponding VAP mutant with 99% PC liposomes as the binding threshold, which is marked by a dotted line and shadow shown in Fig. 3d and Supplementary Fig. 3d,e,g,5e.

##### Calculation of liposome binding coefficient κ_b_

Pairs of raw intensities *F_protein_* and *F_liposome_* were fitted with *F_liposome_* = *κ_b_F_protein_* + *ε* to obtain *κ_b_* and the corresponding r^2^ in MATLAB for each liposome type shown in Fig. 3d and Supplementary Fig. 3f. We only compared *κ_b_* calculated from the same experiment where liposome amounts used can be equalized as described above.

##### κ_b_ calculation and image correction under pH 5

GFP and Rhodamine intensities altered significantly due to pH shift, we thus did intensity correction for a rational *κ_b_* comparison between different pH conditions. We assumed that 1) *κ_b_* of Scs2N-GFP for 99% PC liposomes remained unaffected from pH 7 to pH 5; 2) a linear intensity transformation from pH 7 to pH 5 could apply to both GFP and Rhodamine by introducing simplified correction factors δ_G_ and δ_R_ respectively. Raw intensity data for 99% PC were fitted with 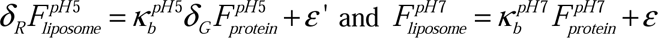 to get δ_G_ and δ_R_, which were used to calculate 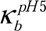 for other liposome type shown in Supplementary Fig. 5a. Original images for pH 5 were correspondingly corrected by δ_G_ or δ_R_ using ImageJ built-in function “Multiply…”.

#### Protein structure modeling

*S. pombe* Pma1 structure (E108-D918) shown in Fig. 6c was modeled using the SWISS-MODEL web server with *S. cerevisiae* Pma1 (7vh5.1.A) ^15^ as the template. Shown intramolecular and intermolecular interactions between the R domain and the FFAT-like motif were predicted according to the autoinhibited structure of *N. crassa* Pma1 (7ny1.1.C)_16._

#### Statistical evaluation

Statistical tests, sample size and definition of error bars and measured values were provided either in figures or figure legends. Boxplots shown in Fig. S1b,S1c,S2d,S2i,3c,S3e,S3g were generated in MATLAB. On each box, the central line indicates the median, and the bottom and top edges of the box indicate the 25th and 75th percentiles, respectively. Whiskers extend to the most extreme data points not considered outliers, and the outliers are plotted individually using the ’+’ symbol. *P*-values and r^2^ were computed using MATLAB ttest2() function and ‘Curve Fitting Toolbox’ respectively. Pearson correlation coefficients (Pearson’s r) in Fig. 2a were calculated in Excel. Cophenetic distances shown in Fig. 2a,6a were calculated from hierarchical clustering using MATLAB and Vector NTI respectively.

## Data availability

The data that support the findings of this study are available from the corresponding author upon reasonable request.

## Acknowledgments

We thank Dr. Yasunori Saheki for sharing the original plasmid to construct GFP-D4H in this study. We are grateful to Dr. Gregory Jedd for sharing human cDNA and protein purification protocol. Authors were funded by the Temasek Life Sciences Laboratory.

## Author Contributions

D.Z. conceived and supervised the project. K.L.H. and D.Z. established liposome assays. K.L.H. and T.S. did protein purification and western blots. B.M. did pH sensitivity tests. B.M., T.S. and K.L.H. performed drug treatment and measured cytosolic pH. B.M., T.S., A.Y.E.N., and A.Q.E.N. did Co-IP experiments. T.S., B.M., K.L.H., A.Y.E.N., and D.Z. performed confocal microscopy. B.M., T.S. and A.Y.E.N. did FRAP analyses. K.L.H., B.M., T.S, D.Z., A.Y.E.N., and A.Q.E.N. constructed strains. D.Z. did interatom, bioinformatics and other quantitative analyses. D.Z. wrote the paper. K.L.H. and B.M. contributed to the manuscript.

## Materials & Correspondence

This study did not generate new unique reagents. Plasmids generated in this study are available upon request. Further information and requests for resources and reagents should be directed to and will be fulfilled by Dan Zhang.

