## Supplementary information for "VAP-mediated membrane tethering mechanisms implicate ER-PM contact function in pH homeostasis"

**Supplementary Figure 1: The MSP** **domain and abundance of VAPs are crucial for keeping the PM PI_4_P level in fission yeast.**

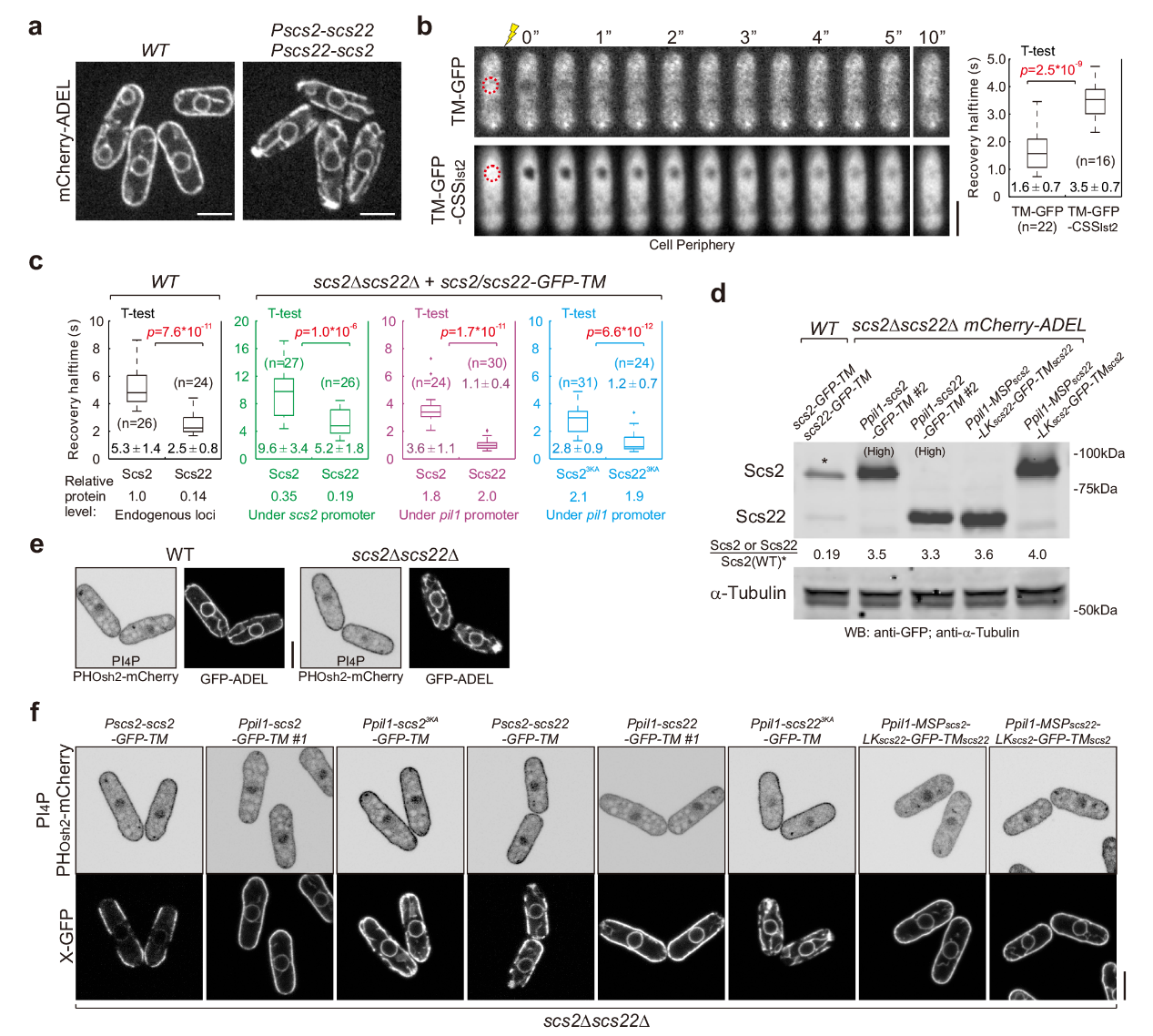

(**a**) Central focal plane spinning disk confocal images of indicated cells expressing mCherry-ADEL. (**b**) Fluorescence recovery after photobleaching (FRAP) analyses of TM-GFP and TM-GFP-CSS_Ist2_ in wild type (*WT*). Shown are top focal planes of representative cells before and after photobleaching, with bleached regions circled. Quantification of the recovery halftime (mean±standard deviation [SD]) of corresponding proteins is shown on the right. (**c**) Quantification of recovery halftime of VAP variants in indicated cells. Relative protein levels are shown as in Figure 1B. n, cell number. *p*-values, two-tailed t test. (**d**) Expression levels of GFP-tagged VAP variants in indicated cells. Relative ratios of protein levels as compared to Scs2 level (asterisked) in *WT* are included. Protein samples were probed with indicated antibodies. (**e**,**f**) Central focal plane scanning confocal images of cells expressing indicated proteins. Scale bars, 5 μm.

**Supplementary Figure 2: Interactome analyses reveal divergent binding preferences between two fission yeast VAPs.**

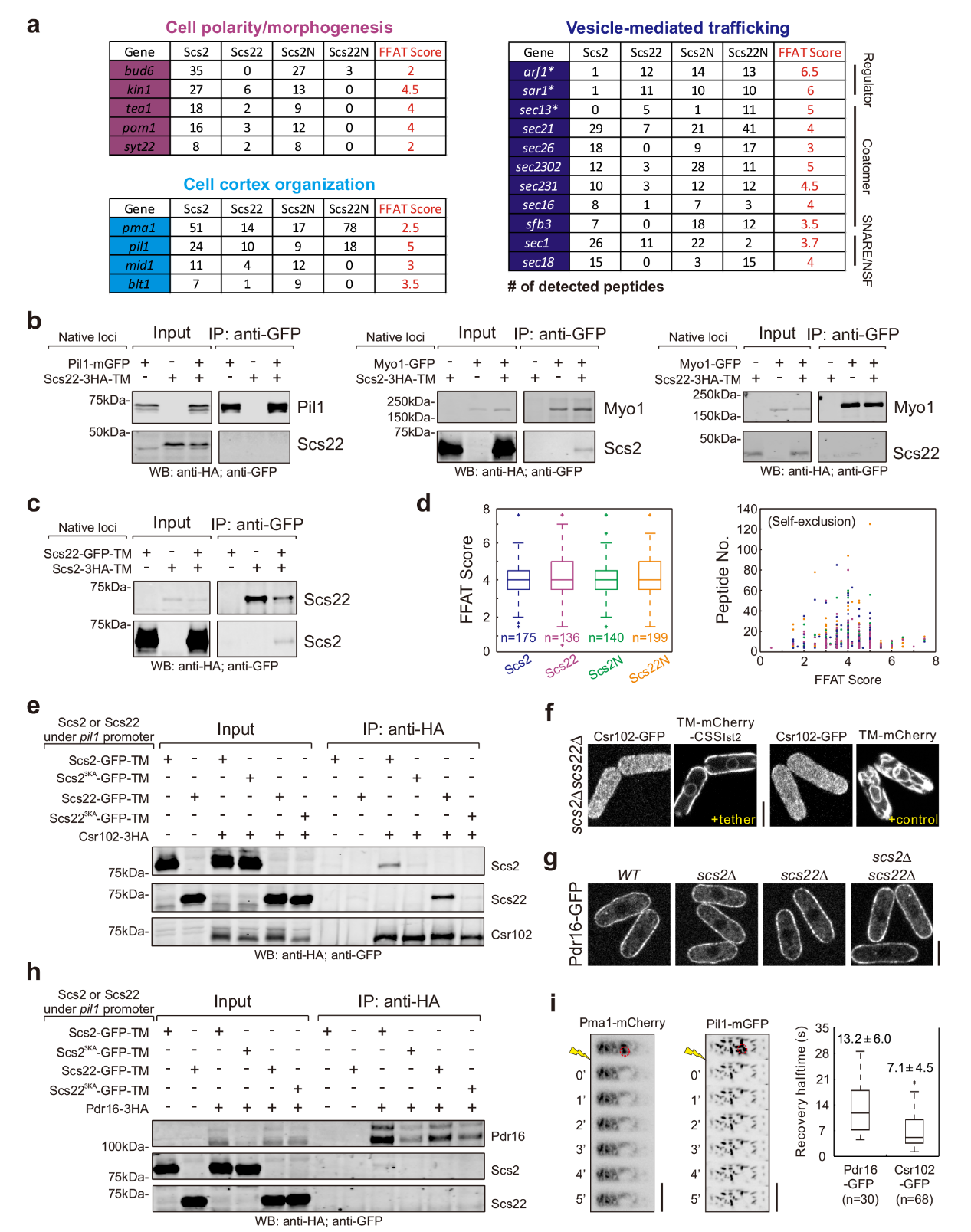

(**a**) Summary of major categories of interactors exhibiting divergent binding preferences towards Scs2 and Scs22. Numbers of detected peptides and the corresponding minimal FFAT scores are included. (**b**,**c**,**e**,**h**) Co-IP of indicated proteins expressed from indicated loci. Samples were probed with indicated antibodies. (**d**) Statistical distribution of minimal FFAT scores of interactors. n, number of total detected interactors for each bait. The minimal FFAT score of each interactor and its corresponding peptide number are plotted on the right. Color indicates the bait. n, number of interactors excluding the bait itself. (**f**,**g**) Central focal plane scanning confocal images of cells expressing indicated proteins. (**i**) FRAP analyses of Pma1-mCherry and Pil1-mGFP in *WT*. Shown are top focal planes of representative cells before and after photobleaching, with bleached regions circled. Quantification of the recovery halftime (mean±SD) of Pdr16-GFP and Csr102-GFP is shown on the right. n, cell number. Scale bars, 5 μm.

**Supplementary Figure 3: The MSP domain is important for the conserved binding of VAPs with major anionic phospholipids.**

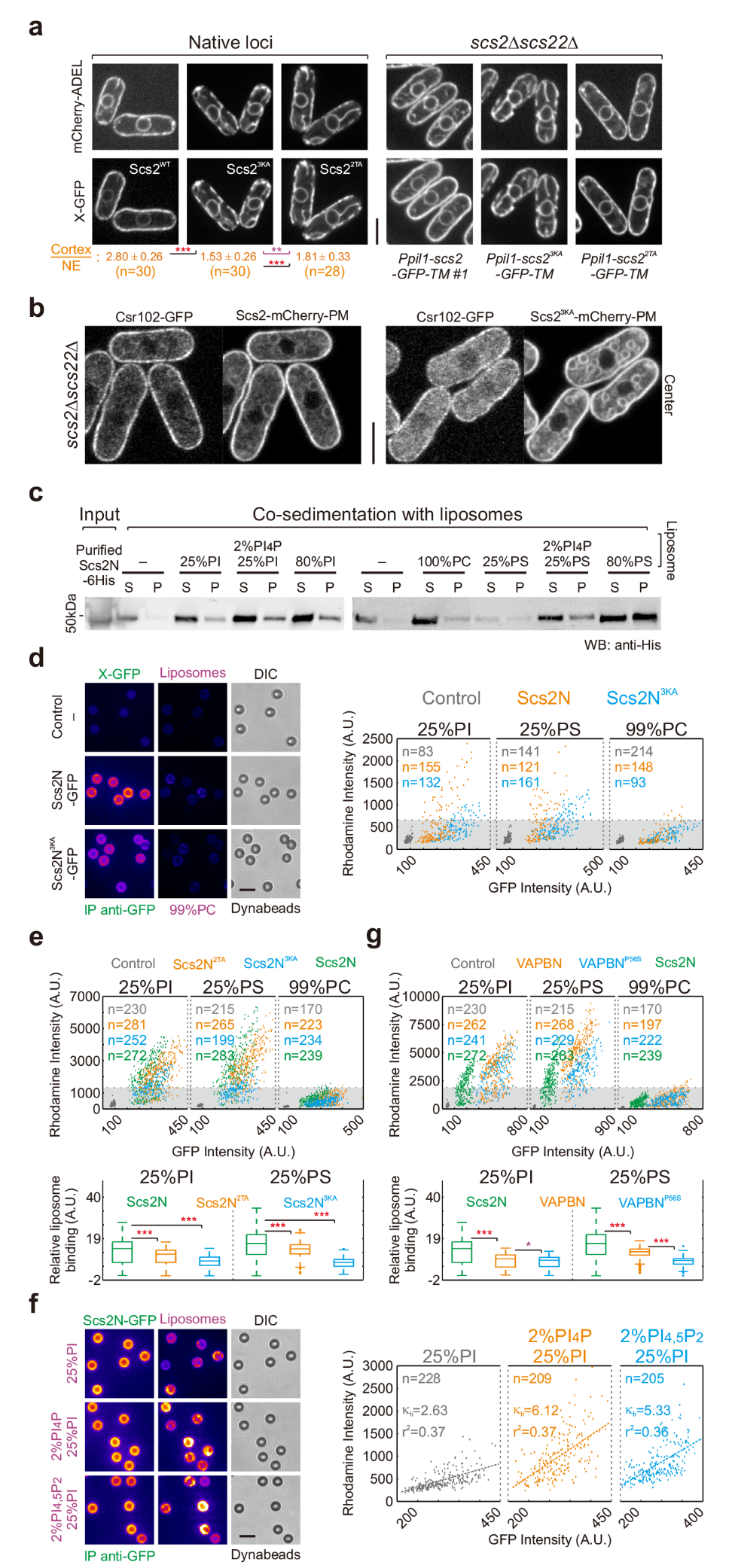

(**a**) Central focal plane spinning disk confocal images of indicated cells expressing indicated proteins. Shown are central focal planes. Normalized cortex/NE ratios (mean±standard deviation [SD]) of Scs2 variants are included. *p*-values, two-tailed t test, ****p*<10^-7^<***p*<0.001; n, cell number. (**b**) Central focal plane scanning confocal images of *scs2Δscs22Δ* cells expressing indicated proteins. (**c**) Liposome co-sedimentation of purified Scs2N-6His. Samples were probed with anti-His antibody. S, supernatant; P, pellet. (**d-g**) Fluorescence microscopy-based protein-liposome binding assays. Indicated GFP-tagged VAP variants immobilized on Dynabeads were incubated with indicated rhodamine-labelled liposomes. In (**d**,**f**), shown are pseudo-colored spinning disk confocal images with the same contrast for each fluorescent channel. In (**e**,**g**), quantifications are shown at bottom panels; *p*-values, two-tailed t test, ****p*<10^-6^<**p*<0.02; Plots from raw intensity data are included in (**d-g**); the dotted line and shadow mark the liposome binding threshold (see Methods for details). κ_b_, liposome binding coefficient; r^2^, r squared; n, bead number. Scale bars, 5 μm.

**Supplementary Figure 4: ER-PM contact formation is compromised in cells lacking inositol and PS.**

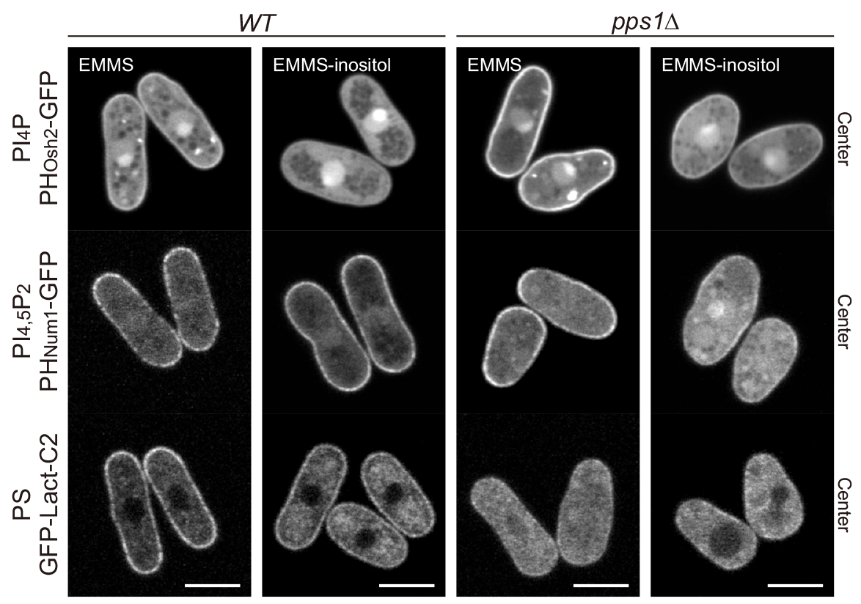

Central focal plane scanning confocal images of indicated cells expressing indicated proteins cultured in the presence or absence of inositol. EMMS, Edinburgh minimal media supplemented with appropriate amino acids. Scale bars, 5 μm.

**Supplementary Figure 5: Binding of Scs2 to PI and PS is pH-sensitive.**

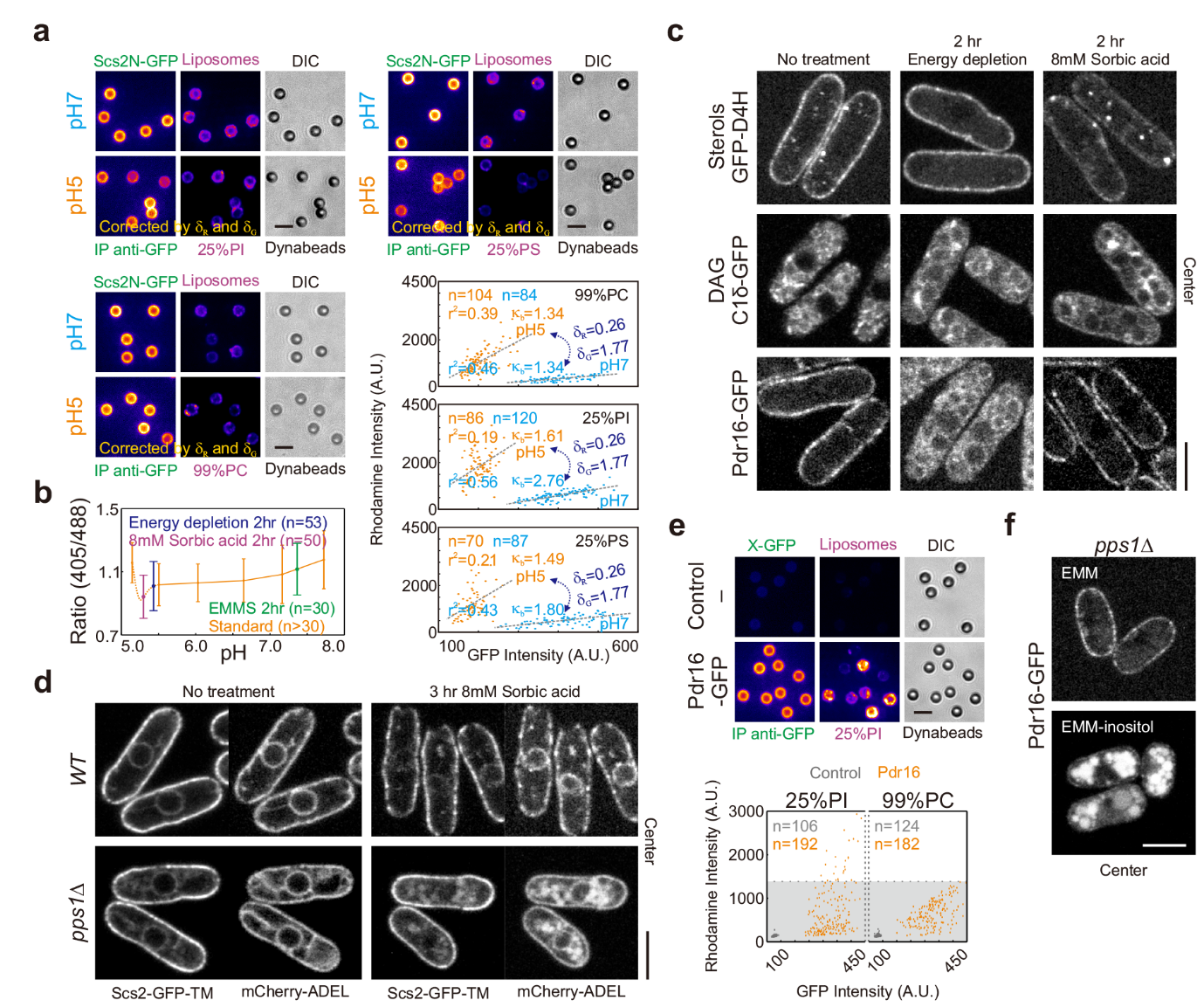

(**a**,**e**) Fluorescence microscopy-based protein-liposome binding assays. GFP-tagged proteins immobilized on Dynabeads were incubated with indicated rhodamine-labelled liposomes. Shown are pseudo-colored spinning disk confocal images with the same contrast for each fluorescent channel. In (**a**), images shown for pH 5 are after correction, by intensity correction factors δ_G_ and δ_R_ respectively for GFP and Rhodamine (see Methods for details). Linear regression plots from raw intensity data are included. In (**e**), the dotted line and shadow mark the liposome binding threshold. n, bead number. κ_b_, liposome binding coefficient. r^2^, r squared. (**b**) Cytosolic pH of sfpHluorin-expressing *WT* cells after various treatments. sfpHluorin standard curve is included. Error bars represent 2×SD. n, cell number. (**c**,**d**,**f**) Central focal plane scanning confocal images of cells expressing indicated proteins after various treatments. Scale bars, 5 μm.

**Supplementary Figure 6: Scs2-interaction is likely important for localizing Pma1-ΔCter to the PM.**

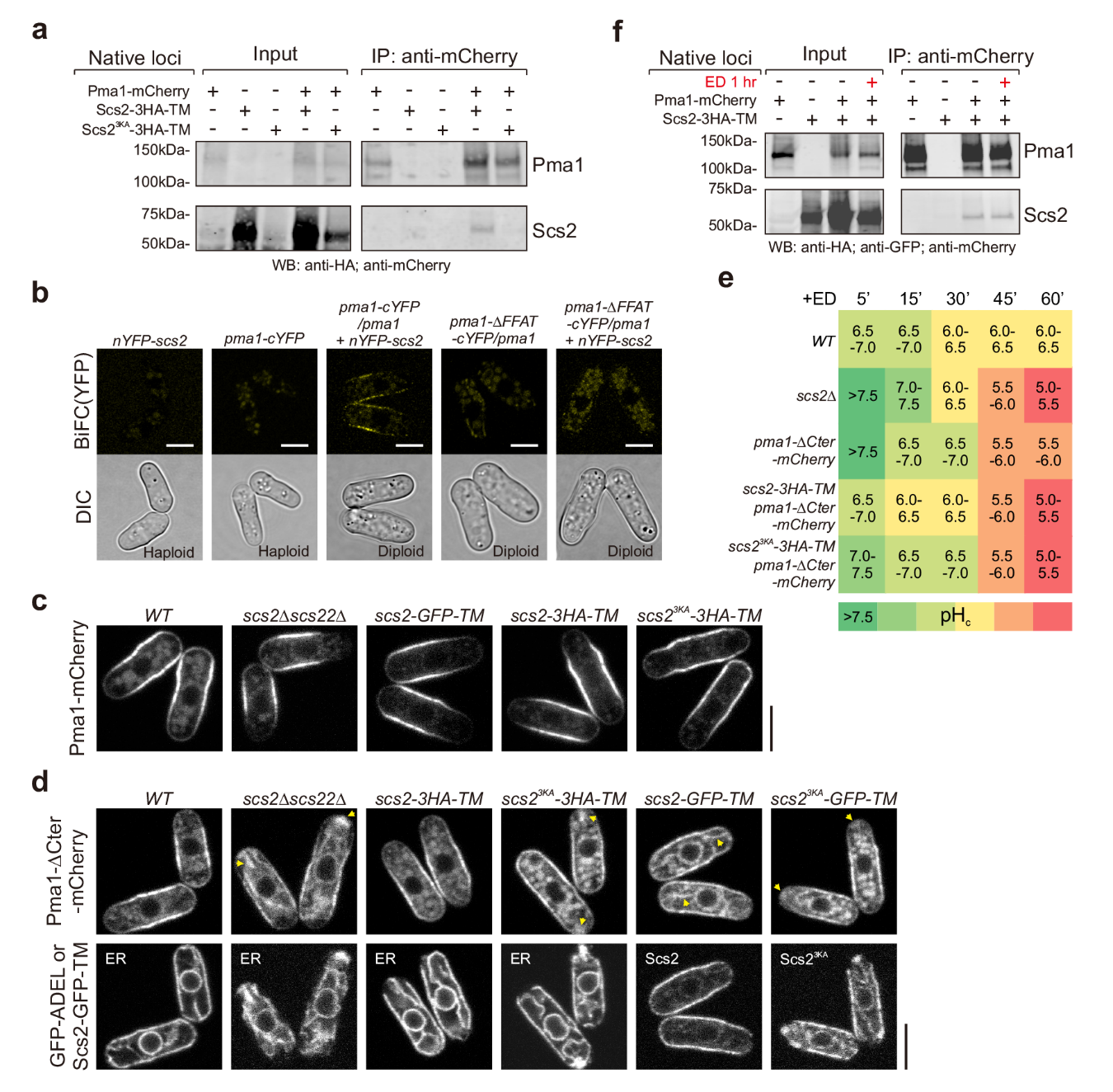

(**a**,**f**) Co-IP of indicated proteins expressed from native loci. Samples were probed with indicated antibodies. (**b**-**d**) Central focal plane scanning confocal images of cells expressing indicated proteins. Yellow arrowheads designate abnormal ER retention of Pma1-ΔCter-mCherry. Scale bars, 5 μm. (**e**) pH_c_ of sfpHluorin-expressing cells with indicated backgrounds at different time points after energy depletion (ED). The times are relative to ED buffer addition. pH_c_ range is indicated by both number and color. For each cell type, n>160 cells were used for measurement.

**Supplementary Table 1: Strain list.**

| **Strain** | **Genotype** | **Figure** | **Reference** |
| --- | --- | --- | --- |
| ZD8 | *scs2Δ::ura4^+^ scs22Δ::ura4^+^ ade6 ura4-D18 leu1-32 h-* | 3c, 3d, S3d, S3e, S3g, 6d | Lab stock |
| ZD12 | *Pnmt1-TM-GFP::leu1^+^ ura4-D18 ade6* | S1b | Lab stock |
| ZD14 | *Pnmt1-TM-GFP-CSS_Ist2_::leu1^+^ ura4-D18 ade6* | S1b | Lab stock |
| ZD45 | *scs2Δ::ura4^+^ scs22Δ::ura4^+^ GFP-ADEL::ura4^+^ leu1-32 ade6* | 1a | Lab stock |
| ZD94 | *scs2-GFP-TM::ura4^+^ ade6-210 ura4-D18 leu1-32 h+* | 2a, 6e | Lab stock |
| ZD106 | *scs2Δ::ura4^+^ GFP-ADEL::leu1^+^ ura4-D18 ade6* | 1a | Lab stock |
| ZD107 | *scs22Δ::ura4^+^ GFP-ADEL::leu1^+^ ura4-D18 ade6* | 1a | Lab stock |
| ZD115 | *sec6-GFP::ura4^+^ mCherry-ADEL::leu1^+^ ura4-D18 ade6 h+* | S1a | Lab stock |
| ZD120 | *scs2-GFP-TM::ura4^+^ mCherry-ADEL::leu1^+^ ura4-D18 ade6* | 1c, S3a, 4b, 5d, S5d | Lab stock |
| ZD204 | *ade6-M210 leu1-32 ura4-D18 h-* | S5e | Lab stock |
| ZD205 | *ade6-M216 leu1-32 ura4-D18 h+* | 6d | Lab stock |
| ZD352 | *GFP-ADEL::leu1^+^ ura4-D18 ade6 h+* | 1a | Lab stock |
| ZD715 | *pma1-mCherry::ura4^+^ ade6 ura4-D18 leu1-32 h+* | 2b, S2i, 6d, 6e, S6a, S6c, S6f | Lab stock |
| ZD746 | *scs2Δ::ura4^+^ scs22Δ::ura4^+^ pma1-mCherry::ura4^+^ ade6 ura4-D18 leu1-32 h+* | 6d | This study |
| ZD973 | *myo1-GFP::kan^+^ ade6 leu1-32 ura4-D18 h+* | S2b | Lab stock |
| ZD1365 | *csr102Δ::kan^+^ sec14Δ::ura4^+^ pdr16Δ::ura4^+^ Prtn1-PH_Osh2_-GFP::ura4^+^ ade6 leu1-32 h-* | 2d | This study |
| ZD1631 | *pdr16Δ::ura4^+^ Prtn1-PH_Osh2_-GFP::ura4^+^ ade6 leu1-32 h+* | 2d | This study |
| ZD1657 | *csr102Δ::ura4^+^ Prtn1-PH_Osh2_-GFP::ura4^+^ ade6 leu1-32 h+* | 2d | This study |
| ZD1667 | *scs2Δ::ura4^+^ scs22Δ::ura4^+^ pdr16-GFP::ura4^+^ ade6-M210 leu1-32 ura4-D18 h-* | S2g | This study |
| ZD1669 | *scs2Δ::ura4^+^ pdr16-GFP::ura4^+^ ade6 leu1-32 ura4-D18 h-* | S2g | This study |
| ZD1671 | *scs22Δ::ura4^+^ pdr16-GFP::ura4^+^ ade6 leu1-32 ura4-D18 h+* | S2g | This study |
| ZD1695 | *scs22-3HA-TM::ura4^+^ ade6-M216 leu1-32 ura4-D18 h+* | 2b, S2b | This study |
| ZD1706 | *pdr16-GFP::ura4^+^ scs2-3HA-TM::ura4^+^ ade6 leu1-32 ura4-D18 h+* | 2b | This study |
| ZD1707 | *csr102-GFP::ura4^+^ scs2-3HA-TM::ura4^+^ ade6 leu1-32 ura4-D18 h+* | 2b | This study |
| ZD1789 | *csr102Δ::ura4^+^ pdr16Δ::ura4^+^ Prtn1-PH_Osh2_-GFP::ura4^+^ ade6 leu1-32 h-* | 2d | This study |
| ZD1949 | *pdr16-GFP::ura4^+^ scs22-3HA-TM::ura4^+^ ade6 leu1-32 ura4-D18 h+* | 2b | This study |
| ZD2054 | *pil1-mGFP::ura4^+^ ade6-M210 leu1-32 ura4-D18 h-* | S2b, S2i | Lab stock |
| ZD2069 | *scs22-3HA-TM::ura4^+^ pil1-mGFP::ura4^+^ ade6 leu1-32 ura4-D18 h+* | S2b | This study |
| ZD2200 | *scs2-3HA-TM::ura4^+^ ade6-M216 leu1-32 ura4-D18 h+* | 2b, S2b, S2c, 6e, S6a, S6f | Lab stock |
| ZD2219 | *csr102-GFP::ura4^+^ scs22-3HA-TM::ura4^+^ ade6 leu1-32 ura4-D18 h+* | 2b | This study |
| ZD2236 | *scs2N(1-360aa)-GFP::ura4^+^ ura4-D18 leu1-32 ade6 h-* | 2a | Lab stock |
| ZD2326 | *pdr16-GFP::ura4^+^ ade6-M210 leu1-32 ura4-D18 h-* | 2b, S2g, S2i, S5c, S5e | This study |
| ZD2405 | *csr102-GFP::ura4^+^ ade6-M210 leu1-32 ura4-D18 h-* | 2b, S2i | This study |
| ZD2406 | *scs22N(1-296aa)-GFP::ura4^+^ ade6-M216 leu1-32 ura4-D18 h-* | 2a | Lab stock |
| ZD2441 | *scs2-GFP-TM::ura4^+^ ade6 leu1-32 ura4-D18 h-* | S1c | Lab stock |
| ZD2445 | *scs22-GFP-TM::ura4^+^ ade6-M210 leu1-32 ura4-D18 h-* | S1c | Lab stock |
| ZD2446 | *scs22-GFP-TM::ura4^+^ ade6-M216 leu1-32 ura4-D18 h+* | 2a, S2c | Lab stock |
| ZD255 | *Prtn1-PH_Osh2_-GFP::ura4^+^ leu1-32 ade6 h-* | 2d | Lab stock |
| ZD2593 | *scs22-GFP-TM::ura4^+^ mCherry-ADEL::ura4^+^ ade6 leu1-32 h+* | 1c | This study |
| ZD2767 | *scs2Δ::Pscs2-scs22::kan^+^ scs22Δ::Pscs22-scs2::kan^+^ mCherry-ADEL::leu1^+^ ade6 ura4-D18 h-* | S1a | This study |
| ZD2846 | *myo1-GFP::kan^+^ scs2-3HA-TM::ura4^+^ ade6 leu1-32 ura4-D18 h+* | S2b | This study |
| ZD2847 | *myo1-GFP::kan^+^ scs22-3HA-TM::ura4^+^ ade6 leu1-32 ura4-D18 h+* | S2b | This study |
| ZD2982 | *sec14Δ::ura4^+^ Prtn1-PH_Osh2_-GFP::ura4^+^ ade6 leu1-32 h-* | 2d | This study |
| ZD3007 | *sec14Δ::ura4^+^ csr102Δ::kan^+^ Prtn1-PH_Osh2_-GFP::ura4^+^ ade6 leu1-32 h-* | 2d | This study |
| ZD3157 | *Prtn1-PH_Num1_-GFP::ura4^+^ ade6 leu1-32 h+* | S4a, 5a | Lab stock |
| ZD3158 | *Prtn1-PH_Osh2_-GFP::ura4^+^ ade6 leu1-32 h+* | S4a, 5a | Lab stock |
| ZD3191 | *sec14Δ::ura4^+^ pdr16Δ::ura4^+^ Prtn1-PH_Osh2_-GFP::ura4^+^ ade6 leu1-32 h+* | 2d | This study |
| ZD3278 | *scs2-GFP-TM::ura4^+^ scs22-GFP-TM::ura4^+^ ade6 leu1-32 ura4-D18 h-* | 1b, S1d | This study |
| ZD3371 | *scs2-3HA-TM::ura4^+^ scs22-GFP-TM::ura4^+^ ade6 leu1-32 ura4-D18 h+* | S2c | This study |
| ZD3413 | *nYFP-scs2::leu1^+^ ade6 ura4-D18 h-* | S6b | This study |
| ZD3428 | *scs2^K36/38/43A^-3HA-TM::kan^+^ ade6 leu1-32 ura4-D18 h-* | S6a | Lab stock |
| ZD3560 | *scs2Δ::ura4^+^ scs22Δ::ura4^+^ Ppil1-scs2-GFP-TM::leu1^+^ mCherry-ADEL::ura4^+^ ade6 #1 h-* | 1b, 1c, S2e, S2h, 3e, S3a | This study |
| ZD3561 | *scs2Δ::ura4^+^ scs22Δ::ura4^+^ Ppil1-scs2-GFP-TM::leu1^+^ mCherry-ADEL::ura4^+^ ade6 #2 h-* | 1e, S1c, S1d | This study |
| ZD3565 | *scs2Δ::ura4^+^ scs22Δ::ura4^+^ Ppil1-scs22-GFP-TM::leu1^+^ mCherry-ADEL::ura4^+^ #1 ade6 h-* | 1b, 1c, S1c, S2e, S2h | This study |
| ZD3566 | *scs2Δ::ura4^+^ scs22Δ::ura4^+^ Ppil1-scs22-GFP-TM::leu1^+^ mCherry-ADEL::ura4^+^ ade6 #2 h-* | 1e, S1d | This study |
| ZD3577 | *csr102Δ::ura4^+^ pdr16Δ::ura4^+^ pil1Δ::ura4^+^ pma1-mCherry::ura4^+^ scs2-GFP-TM::ura4^+^ ade6 leu1-32 ura4-D18 h-* | 2g | This study |
| ZD3608 | *scs2Δ::ura4^+^ scs22Δ::ura4^+^ Pscs2-scs2-GFP-TM::leu1^+^ mCherry-ADEL::ura4^+^ ade6 h-* | 1b, 1c, S1c | This study |
| ZD3611 | *scs2Δ::ura4^+^ scs22Δ::ura4^+^ Pscs2-scs22-GFP-TM::leu1^+^ mCherry-ADEL::ura4^+^ ade6 h+* | 1b, 1c, S1c | This study |
| ZD3647 | *scs2Δ ::ura4^+^ scs22Δ::ura4^+^ Ppil1-scs2^K36/38/43A^-GFP-TM::leu1^+^ mCherry-ADEL::ura4^+^ ade6 h-* | 1b, 1c, S1c, S3a, 5d | This study |
| ZD3676 | *csr102Δ::ura4^+^ pdr16Δ::ura4^+^ pil1Δ::ura4^+^ GFP-ADEL::leu1^+^ ade6 ura4-D18 h+* | 2g | This study |
| ZD3756 | *Ppil1-nGFP-D4H::leu1^+^ ura4-D18 ade6 h+* | S5c | This study |
| ZD3898 | *ade6^+^::Pact1-nGFP-Lact-C2::hyg^+^ leu1-32 ura4-D18 h+* | S4a, 5a | This study |
| ZD4000 | *csr102Δ::ura4^+^ pdr16Δ::ura4^+^ pil1Δ::ura4^+^ scs2-GFP-TM::ura4^+^ ade6 leu1-32 ura4-D18 h-* | 2g | This study |
| ZD4005 | *pma1-mCherry::ura4^+^ scs2-GFP-TM::ura4^+^ ade6 leu1-32 ura4-D18 h-* | 6d, 6e, S6c | This study |
| ZD4031 | *pma1-mCherry::ura4^+^ scs2-3HA-TM::ura4^+^ ade6 leu1-32 ura4-D18 h-* | 2b, 6d, 6e, S6a, S6c, S6f | This study |
| ZD4032 | *pma1-mCherry::ura4^+^ scs22-3HA-TM::ura4^+^ ade6 leu1-32 ura4-D18 h-* | 2b | This study |
| ZD4129 | *scs2Δ::ura4^+^ scs22Δ::ura4^+^ mCherry-ADEL::ura4^+^ Ppil1-MSP_scs22_-LK_scs2_-GFP-TM_scs22_ ::leu1^+^ ade6 h-* | 1e, S1d | This study |
| ZD4132 | *scs2Δ::ura4^+^ scs22Δ::ura4^+^ mCherry-ADEL::ura4^+^ Ppil1-MSP_scs2_-LK_scs22_-GFP-TM_scs2_ ::leu1^+^ ade6 h-* | 1e, S1d | This study |
| ZD4133 | *scs2Δ::ura4^+^ scs22Δ::ura4^+^ Ppil1-scs22^K36/38/43A^-GFP-TM::leu1^+^ mCherry-ADEL::ura4^+^ ade6 h-* | 1b, 1c, S1c | This study |
| ZD4238 | *pma1-ΔFFAT-3HA::ura4^+^/pma1 leu1-32/leu1-32 ura4-D18/ura4-D18* | 6a | This study |
| ZD4239 | *pma1-FFAT_Opi1_-3HA::ura4^+^/pma1 leu1-32/leu1-32 ura4-D18/ura4-D18* | 6a | This study |
| ZD4243 | *pma1-4L-3HA::ura4^+^/pma1 leu1-32/leu1-32 ura4-D18/ura4-D18* | 6a | This study |
| ZD4312 | *pma1-cYFP::ura4^+^/pma1 nYFP-scs2::leu1^+^/nYFP-scs2::leu1^+^ ade6/ade6 h-* | S6b | This study |
| ZD4397 | *scs2Δ::ura4^+^ scs22Δ::ura4^+^ Ppil1-scs2^K36/38/43A^-GFP-TM::leu1^+^ csr102-mCherry::ura4^+^ ade6 ura4-D18 h-* | 2f | This study |
| ZD4398 | *scs2Δ::ura4^+^ scs22Δ::ura4^+^ Ppil1-scs2-GFP-TM::leu1^+^ csr102-mCherry::ura4^+^ ade6 ura4-D18 h+* | 2f | This study |
| ZD4399 | *scs2Δ::ura4^+^ scs22Δ::ura4^+^ Ppil1-scs22^K36/38/43A^-GFP-TM::leu1^+^ csr102-mCherry::ura4^+^ ade6 ura4-D18 h-* | 2f | This study |
| ZD4400 | *scs2Δ::ura4^+^ scs22Δ::ura4^+^ Ppil1-scs2^K36/38/43A^-GFP-TM::leu1^+^ mCherry-ADEL::ura4^+^ csr102-3HA::ura4^+^ ade6 h-* | S2e | This study |
| ZD4401 | *scs2Δ::ura4^+^ scs22Δ::ura4^+^ Ppil1-scs2-GFP-TM::leu1^+^ mCherry-ADEL::ura4^+^ csr102-3HA::ura4^+^ ade6 h-* | S2e | This study |
| ZD4449 | *scs2Δ::ura4^+^ scs22Δ::ura4^+^ Ppil1-scs22-GFP-TM::leu1^+^ csr102-mCherry::ura4^+^ ade6 ura4-D18 h-* | 2f | This study |
| ZD4480 | *scs2Δ::ura4^+^ scs22Δ::ura4^+^ Pscs2-scs2-GFP-TM::leu1^+^ csr102-mCherry::ura4^+^ ade6 ura4-D18 h-* | 2f | This study |
| ZD4481 | *scs2Δ::ura4^+^ scs22Δ::ura4^+^ Ppil1-scs22^K36/38/43A^-GFP-TM::leu1^+^ mCherry-ADEL::ura4^+^ csr102-3HA::kan^+^ ade6 h-* | S2e | This study |
| ZD4482 | *scs2Δ::ura4^+^ scs22Δ::ura4^+^ Ppil1-scs22-GFP-TM::leu1^+^ mCherry-ADEL::ura4^+^ pdr16-3HA::kan^+^ ade6 h-* | S2h | This study |
| ZD4483 | *scs2Δ::ura4^+^ scs22Δ::ura4^+^ Ppil1-scs22^K36/38/43A^-GFP-TM::leu1^+^ mCherry-ADEL::ura4^+^ pdr16-3HA::kan^+^ ade6 h-* | S2h | This study |
| ZD4487 | *csr102-GFP::ura4^+^ mCherry-ADEL::leu1^+^ ade6 ura4-D18 h-* | 2e | This study |
| ZD4488 | *scs2Δ::ura4^+^ csr102-GFP::ura4^+^ mCherry-ADEL::leu1^+^ ade6 ura4-D18 h+* | 2e | This study |
| ZD4489 | *scs22Δ::ura4^+^ csr102-GFP::ura4^+^ mCherry-ADEL::leu1^+^ ade6 ura4-D18 h-* | 2e | This study |
| ZD4490 | *scs2Δ::ura4^+^ scs22Δ::ura4^+^ csr102-GFP::ura4^+^ mCherry-ADEL::leu1^+^ ade6 ura4-D18 h-* | 2e | This study |
| ZD4497 | *scs2Δ::ura4^+^ scs22Δ::ura4^+^ csr102-GFP::ura4^+^ Pnmt1-TM-mCherry-CSS_Ist2_::leu1^+^ ade6 ura4-D18 h-* | S2f | This study |
| ZD4498 | *scs2Δ::ura4^+^ scs22Δ::ura4^+^ csr102-GFP::ura4^+^ Pnmt1-TM-mCherry::leu1^+^ ade6 ura4-D18 h+* | S2f | This study |
| ZD4501 | *scs2Δ::ura4^+^ scs22Δ::ura4^+^ Ppil1-scs2^K36/38/43A^-GFP-TM::leu1^+^ mCherry-ADEL::ura4^+^ pdr16-3HA::kan^+^ ade6 h-* | S2h | This study |
| ZD4502 | *scs2Δ::ura4^+^ scs22Δ::ura4^+^ Ppil1-scs2-GFP-TM::leu1^+^ mCherry-ADEL::ura4^+^ pdr16-3HA::kan^+^ ade6 h-* | S2h | This study |
| ZD4503 | *scs2Δ::ura4^+^ scs22Δ::ura4^+^ Ppil1-scs22-GFP-TM::leu1^+^ mCherry-ADEL::ura4^+^ csr102-3HA::kan^+^ ade6 h-* | S2e | This study |
| ZD4517 | *scs2Δ::ura4^+^ scs22Δ::ura4^+^ Pscs2-scs22-GFP-TM::leu1^+^ csr102-mCherry::ura4^+^ ade6 ura4-D18 h+* | 2f | This study |
| ZD4543 | *Prtn1-PH_Osh2_-mCherry::ura4^+^ GFP-ADEL::leu1^+^ ade6 h+* | S1e | This study |
| ZD4544 | *scs2Δ::ura4^+^ scs22Δ::ura4^+^ Prtn1-PH_Osh2_-mCherry::ura4^+^ GFP-ADEL::leu1^+^ ade6 h-* | S1e | This study |
| ZD4546 | *scs2Δ::ura4^+^ scs22Δ::ura4^+^ Ppil1-scs2-GFP-TM::leu1^+^ Prtn1-PH_Osh2_-mCherry::ura4^+^ ade6 h-* | S1f | This study |
| ZD4558 | *scs2Δ::ura4^+^ scs22Δ::ura4^+^ Ppil1-scs22-GFP-TM::leu1^+^ Prtn1-PH_Osh2_-mCherry::ura4^+^ ade6 h-* | S1f | This study |
| ZD4559 | *scs2Δ::ura4^+^ scs22Δ::ura4^+^ Pscs2-scs2-GFP-TM::leu1^+^ Prtn1-PH_Osh2_-mCherry::ura4^+^ ade6 h-* | S1f | This study |
| ZD4560 | *scs2Δ::ura4^+^ scs22Δ::ura4^+^ Ppil1-scs2^K36/38/43A^-GFP-TM::leu1^+^ Prtn1-PH_Osh2_-mCherry::ura4^+^ ade6 h-* | S1f | This study |
| ZD4561 | *scs2Δ::ura4^+^ scs22Δ::ura4^+^ Pscs2-scs22-GFP-TM::leu1^+^ Prtn1-PH_Osh2_-mCherry::ura4^+^ ade6 h-* | S1f | This study |
| ZD4562 | *scs2Δ::ura4^+^ scs22Δ::ura4^+^ Prtn1-PH_Osh2_-mCherry::ura4^+^ Ppil1-MSP_scs2_-LK_scs22_-GFP-TM_scs2_ ::leu1^+^ ade6 h-* | S1f | This study |
| ZD4563 | *scs2Δ::ura4^+^ scs22Δ::ura4^+^ Ppil1-scs22^K36/38/43A^-GFP-TM::leu1^+^ Prtn1-PH_Osh2_-mCherry::ura4^+^ ade6 h-* | S1f | This study |
| ZD4564 | *pma1-ΔFFAT-cYFP::ura4^+^/pma1 ade6/ade6 leu1-32/leu1-32* | S6b | This study |
| ZD4569 | *scs2Δ::ura4^+^ scs22Δ::ura4^+^ Prtn1-PH_Osh2_-mCherry::ura4^+^ Ppil1-MSP_scs22_-LK_scs2_-GFP-TM_scs22_ ::leu1^+^ ade6 h-* | S1f | This study |
| ZD4638 | *pma1-3HA::ura4^+^/pma1 leu1-32/leu1-32 ura4-D18/ura4-D18* | 6a | This study |
| ZD4700 | *scs2Δ::ura4^+^ scs22Δ::ura4^+^ GFP-ADEL::ura4^+^ Pnmt1-scs2-mCherry-PM::leu1^+^ ade6 h-* | 3a, 4c | This study |
| ZD4701 | *scs2Δ::ura4^+^ scs22Δ::ura4^+^ GFP-ADEL::ura4^+^ Pnmt1-scs2^K36/38/43A^-mCherry-PM::leu1^+^ ade6 h-* | 3a | This study |
| ZD4710 | *scs2Δ::ura4^+^ scs22Δ::ura4^+^ Prtn1-PH_Osh2_-GFP::ura4^+^ Pnmt1-scs2-mCherry-PM::leu1^+^ ade6 h-* | 3b | This study |
| ZD4717 | *scs2Δ::ura4^+^ scs22Δ::ura4^+^ Prtn1-PH_Osh2_-GFP::ura4^+^ Pnmt1-scs2^K36/38/43A^-mCherry-PM::leu1^+^ ade6 h-* | 3b | This study |
| ZD4853 | *scs2Δ::ura4^+^ scs22Δ::ura4^+^ csr102-GFP::ura4^+^ Ppil1-scs2-mCherry-PM::leu1^+^ ade6 ura4-D18 h-* | S3b | This study |
| ZD4854 | *scs2Δ::ura4^+^ scs22Δ::ura4^+^ csr102-GFP::ura4^+^ Ppil1-scs2^K36/38/43A^-mCherry-PM::leu1^+^ ade6 ura4-D18 h+* | S3b | This study |
| ZD5096 | *Pscs2-C1δ-GFP::ura4^+^ ade6-704 leu1-32 h-* | S5c | This study |
| ZD5169 | *pps1Δ::kan^+^ scs2Δ::ura4^+^ scs22Δ::ura4^+^ GFP-ADEL::ura4^+^ Pnmt1-scs2-mCherry-PM::leu1^+^ ade6 h-* | 4c | This study |
| ZD5185 | *pdr16-GFP::ura4^+^ scs2-mCherry-TM::kan^+^ ade6 leu1-32 ura4-D18 h+* | 5d | This study |
| ZD5188 | *pps1Δ::kan^+^ Prtn1-PH_Osh2_-GFP::ura4^+^ ade6 leu1-32 h+* | S4a | This study |
| ZD5189 | *pps1Δ::kan^+^ Prtn1-PH_Num1_-GFP::ura4^+^ ade6 leu1-32* | S4a | This study |
| ZD5195 | *pps1Δ::kan^+^ scs2-GFP-TM::ura4^+^ mCherry-ADEL::leu1^+^ ade6 ura4-D18 h-* | 4b, S5d, Movie S1 | This study |
| ZD5209 | *mCherry-ADEL::ura4^+^ Ppil1-sfpHluorin::leu1^+^ ade6 h+* | 5b, Movie S2 | This study |
| ZD5210 | *Ppil1-sfpHluorin::leu1^+^ ade6-M210 ura4-D18 h-* | 6f, S6e | This study |
| ZD5212 | *scs2-mCherry-TM::kan^+^ Ppil1-sfpHluorin::leu1^+^ ade6 leu1-32 h+* | 5c, S5b | This study |
| ZD5293 | *ade6^+^::Pact1-nGFP-Lact-C2::hyg^+^ scs2-mCherry-TM::ura4^+^ leu1-32 ura4-D18 h+* | 5d | This study |
| ZD5295 | *pps1Δ::kan^+^ ade6^+^::Pact1-nGFP-Lact-C2::hyg^+^ leu1-32 ura4-D18 h+* | S4a | This study |
| ZD5296 | *pps1Δ::kan^+^ pdr16-GFP::ura4^+^ ade6 leu1-32 ura4-D18 h-* | S5f | This study |
| ZD5311 | *Prtn1-PH_Osh2_-GFP::ura4^+^ scs2-mCherry-TM::ura4^+^ ade6 leu1-32 h+* | 5d | This study |
| ZD5334 | *Prtn1-PH_Num1_-GFP::ura4^+^ scs2-mCherry-TM::ura4^+^ ade6 leu1-32 h+* | 5d | This study |
| ZD5400 | *scs2Δ::ura4^+^ scs22Δ::ura4^+^ Ppil1-scs2N^K36/K38/K43A^(1-360aa)-GFP::leu1^+^ ade6 ura4-D18 h-* | 3c, S3d, S3e | This study |
| ZD5402 | *pma1-ΔCter(897-919aa)-3HA::ura4^+^ ade6-M216 leu1-32 ura4-D18 h+* | 6d | This study |
| ZD5417 | *scs2Δ::ura4^+^ scs22Δ::ura4^+^ Ppil1-scs2N(1-360aa)-GFP::leu1^+^ ade6 ura4-D18 h-* | 3c, 3d, S3d, S3e, S3f, S3g, S5a | This study |
| ZD5420 | *scs2Δ::ura4^+^ pma1-ΔCter(897-919aa)-3HA::ura4^+^ ade6 leu1-32 ura4-D18 h+* | 6d | This study |
| ZD5421 | *scs22Δ::ura4^+^ pma1-ΔCter(897-919aa)-3HA::ura4^+^ ade6 leu1-32 ura4-D18 h-* | 6d | This study |
| ZD5422 | *scs2Δ::ura4^+^ scs22Δ::ura4^+^ pma1-ΔCter(897-919aa)-3HA::ura4^+^ ade6 leu1-32 ura4-D18 h-* | 6d | This study |
| ZD5482 | *scs2Δ::ura4^+^ scs22Δ::ura4^+^ Ppil1-VAPBN(1-213aa)-GFP::leu1^+^ ade6 ura4-D18 h+* | 3d, S3g | This study |
| ZD5509 | *scs2Δ::ura4^+^ scs22Δ::ura4^+^ mCherry-ADEL::ura4^+^ Ppil1-VAPB-GFP-TM::leu1^+^ ade6 h-* | 3e | This study |
| ZD5510 | *scs2Δ::ura4^+^ scs22Δ::ura4^+^ Ppil1-VAPBN^P56S^ (1-213aa)-GFP::leu1^+^ ade6 ura4-D18 h+* | 3d, S3g | This study |
| ZD5512 | *scs2Δ::ura4^+^ scs22Δ::ura4^+^ Ppil1-scs22N(1-296aa)-GFP::leu1+ ade6 h-* | 3d | This study |
| ZD5513 | *scs2Δ::ura4^+^ scs22Δ::ura4^+^ Ppil1-scs22N^K36/K38/K43A^(1-296aa)-GFP::leu1+ ade6 h-* | 3d | This study |
| ZD5520 | *scs2Δ::ura4^+^ scs22Δ::ura4^+^ mCherry-ADEL::ura4^+^ Ppil1-VAPB^P56S^-GFP-TM::leu1^+^ ade6 h-* | 3e | This study |
| ZD5537 | *scs2^K36/38/43A^-3HA-TM::kan^+^ pma1-mCherry::ura4^+^ ade6 leu1-32 ura4-D18* | 6d, S6a, S6c | This study |
| ZD5553 | *pma1-ΔCter(897-919aa)-mCherry::ura4^+^ ade6-M216 leu1-32 ura4-D18 h+* | 6d | This study |
| ZD5561 | *scs2-3HA-TM::ura4^+^ pma1-ΔCter(897-919aa)-mCherry::ura4^+^ ade6 leu1-32 ura4-D18* | 6d | This study |
| ZD5562 | *scs2-GFP-TM::ura4^+^ pma1-ΔCter(897-919aa)-mCherry::ura4^+^ ade6 leu1-32 ura4-D18 h-* | 6d, S6d | This study |
| ZD5563 | *scs2^K36/38/43A^-3HA-TM::kan^+^ pma1-ΔCter(897-919aa)-mCherry::ura4^+^ ade6 leu1-32 ura4-D18 h-* | 6d | This study |
| ZD5564 | *scs2^K36/38/43A^-GFP-TM::kan^+^ pma1-ΔCter(897-919aa)-mCherry::ura4^+^ ade6 leu1-32 ura4-D18* | 6d, S6d | This study |
| ZD5598 | *scs2Δ::ura4^+^ Ppil1-sfpHluorin::leu1^+^ ade6 ura4-D18 h-* | S6e | This study |
| ZD5601 | *scs2^K36/38/43A^-3HA-TM::kan^+^ pma1-ΔCter(897-919aa)-mCherry::ura4^+^ Ppil1-sfpHluorin::leu1^+^ ade6 ura4-D18* | S6e | This study |
| ZD5606 | *pma1-ΔCter(897-919aa)-mCherry::ura4^+^ Ppil1-sfpHluorin::leu1^+^ ade6 ura4-D18 h+* | S6e | This study |
| ZD5607 | *scs2-3HA-TM::ura4^+^ pma1-ΔCter(897-919aa)-mCherry::ura4^+^ Ppil1-sfpHluorin::leu1^+^ ade6 ura4-D18* | S6e | This study |
| ZD5620 | *scs2^K36/38/43A^-3HA-TM::kan^+^ pma1-ΔCter(897-919aa)-mCherry::ura4^+^ GFP-ADEL::leu1^+^ ade6 ura4-D18 h-* | S6d | This study |
| ZD5624 | *pma1-mCherry::ura4^+^ GFP-ADEL::leu1^+^ ade6 ura4-D18 h+* | S6d | This study |
| ZD5625 | *scs2Δ::ura4^+^ scs22Δ::ura4^+^ pma1-mCherry::ura4^+^ GFP-ADEL::leu1^+^ ade6 ura4-D18 h+* | S6c | This study |
| ZD5628 | *scs2Δ::ura4^+^ scs22Δ::ura4^+^ pma1-ΔCter(897-919aa)-mCherry::ura4^+^ GFP-ADEL::leu1^+^ ade6 ura4-D18 h-* | S6d | This study |
| ZD5631 | *scs2-3HA-TM::ura4^+^ pma1-ΔCter(897-919aa)-mCherry::ura4^+^ GFP-ADEL::leu1^+^ ade6 ura4-D18 h-* | S6d | This study |
| ZD5740 | *nYFP-scs2::leu1^+^/nYFP-scs2::leu1^+^ pma1-ΔFFAT-cYFP::ura4^+^/pma1 ade6 h-* | S6b | This study |
| ZD5744 | *scs2-GST-TM::ura4^+^ pma1-mCherry::ura4^+^ ade6 leu1-32 h-* | 6d, 6e | This study |
| ZD5745 | *scs2-GST-TM::ura4^+^ pma1-mCherry::ura4^+^ ade6 leu1-32 h-* | 6d | This study |
| ZD5763 | *scs2-GST-TM::ura4^+^ pma1-mCherry::ura4^+^ his5^+^::Ppil1-sfpHluorin::bsd^+^ ade6 leu1-32 h-* | 6f | This study |
| ZD5764 | *scs2-GST-TM::ura4^+^ pma1-mCherry::ura4^+^ his5^+^::Ppil1-sfpHluorin::bsd^+^ ade6 leu1-32 h-* | 6f | This study |
| ZD5821 | *scs2-GST-TM::ura4^+^ ade6-M210 leu1-32 h-* | 6e | This study |
| ZD5962 | *pma1-mCherry::ura4^+^ scs2-3HA-TM::ura4^+^ his5^+^::Ppil1-sfpHluorin::bsd^+^ ade6 leu1-32 h-* | 6f | This study |
| ZD5965 | *pma1-cYFP::ura4^+^ ade6 leu1-32 h+* | S6b | This study |
| ZD5985 | *scs2Δ::ura4^+^ scs22Δ::ura4^+^ mCherry-ADEL::ura4^+^ Ppil1-scs2^T39/T40A^-GFP-TM::leu1^+^ ade6 h-* | S3a | This study |
| ZD5986 | *scs2Δ::ura4^+^ scs22Δ::ura4^+^ Ppil1-scs2N^T39A/T40A^(1-360aa)-GFP::leu1^+^ ade6 h-* | S3e | This study |
| ZD6031 | *kan^+^::scs2-cYFP-TM ade6 leu1-32 h-* | 6b | This study |
| ZD6043 | *scs2^K36A/K38A/K43A^-GFP-TM::kan^+^ Pbip1-mCherry-ADEL::ura4^+^ ade6-704 leu1-32 h-* | S3a | This study |
| ZD6044 | *scs2N^T39/T40A^(1-360aa)-GFP-TM::kan^+^ Pbip1-mCherry-ADEL::ura4^+^ ade6-704 leu1-32 h-* | S3a | This study |
| ZD6053 | *pma1-nYFP::ura4^+^ ade6-M216 leu1-32 h+* | 6b | This study |
| ZD6085 | *kan^+^::scs2-cYFP-TM pma1-nYFP::ura4^+^ ade6 leu1-32 h+* | 6b | This study |

**Supplementary Movie Legends**

**Supplementary Movie 1.** 3D Rendering of Scs2-GFP-TM-expressing *WT* and *pps1Δ* cells after inositol starvation. The movie (at 15 fps) refers to Fig. 4b.

**Supplementary Movie 2.** 3D Rendering of mCherry-ADEL-expressing *WT* cells after indicated treatments. The movie (at 15 fps) refers to Fig. 5b.
